# A dual neural system supporting multi-tasking cognitive flexibility

**DOI:** 10.64898/2026.01.27.702157

**Authors:** Mengya Zhang, Qing Yu

## Abstract

One ubiquitous feature of human intelligence is the ability to flexibly switch between multiple tasks. Abstract task representations provide a basis for task learning, switching, and generalization, yet how the brain coordinates multiple task representations to support multitasking and task-switching remains poorly understood. We recorded functional MRI activity in human participants while they concurrently held multiple task rules in working memory and prioritized different rules for stimulus processing across trials. Our results reveal two distinct coding schemes for task prioritization. First, an active coding scheme supports the spatial separation of prioritized and unprioritized task rules: prioritized rules are represented across a distributed cortical network, whereas unprioritized rules are only found in the posterior cortex. Second, besides this active scheme, subregions of the default mode network, including the medial prefrontal cortex and hippocampus, contribute to offloading task representations into an unprioritized state and subsequently maintain sustained representations of these rules across trials via latent neural codes. Behavioral predictions using on-task, trial-wise and sustained, block-wise representations further support the neural dissociation. These findings unveil a dual neural system with distinct coding schemes that jointly enable cognitive flexibility for task implementation and switching.

## Introduction

In everyday life, individuals often need to perform multiple tasks within limited time windows - for instance, picking up a food order, refilling gas, and doing grocery shopping before driving back home. A common strategy for managing such demands is to perform tasks sequentially, that is, prioritizing one task while putting others on hold for later execution. Successfully implementing this strategy requires the ability to concurrently maintain multiple tasks, to differentiate between them based on their priority state, and to switch between them flexibly when needed. Despite its critical role in everyday cognition, the underlying neural mechanisms supporting this cognitive flexibility remain largely unknown.

Flexible and efficient task-switching relies on the formation of task representations, that is, abstract representations of task rules, structures, or schemas that guide context-dependent stimulus-action mappings ^1–4^. Having such abstract representations reduces the need for incremental learning (i.e., re-learning each task from scratch), thereby allowing individuals to execute tasks more effectively and update them flexibly in working memory (WM) ^5^. Although task representations have been studied extensively in both cognitive control ^6^ and cognitive map^7, 8^ research, surprisingly few studies have investigated how multiple task representations are coordinated within WM to support on-demand task prioritization and deprioritization. This gap highlights the need for an experimental paradigm that can directly probe these processes.

Research on stimulus prioritization in WM provides a useful framework for addressing this question ^9–12^. Here, stimulus refers to a concrete mnemonic unit with feature-specific content, such as visual colors or objects, as distinct from abstract task information. When multiple stimuli are maintained in WM, the item that will be immediately tested is considered prioritized, whereas the ones to be tested later are unprioritized. Previous studies have demonstrated that prioritized and unprioritized stimuli are maintained through distinct neural mechanisms. The most prominent finding is that the prioritized stimulus is actively represented by a distributed WM network, including frontal, parietal, and sensory regions ^13^, all of which have been implicated in WM-related processes ^14–16^. This active representation is typically characterized by stimulus-specific delay-period activity, a core neural signature of WM ^17, 18^, including persistent neural activity at the single-neuron level, and successful population-level decoding of stimulus information using human fMRI ^15, 16, 19^.

In contrast to active representation of prioritized information during WM, the unprioritized stimulus often exhibits weaker or even no significant representation in these regions ^20–22^, suggesting that the maintenance of unprioritized information may involve distinct neural mechanisms, possibly through latent neural codes. Accordingly, several theoretical accounts have been proposed for the neural coding of unprioritized information. The “activity-silent” account proposes that instead of being stored in active spiking activity, unprioritized information is silently encoded in short-term synaptic plasticity that cannot be detected at the activity level ^23–25^. The recoded representation account suggests that unprioritized information is also actively represented, but in a format rotated or recoded relative to the prioritized information to reduce interference ^26–28^. A third account, the spatial separation account, posits that unprioritized stimulus is (weakly) represented in restricted brain regions, such as frontoparietal regions, rather than in widespread networks as the prioritized stimulus ^13^. Notably, these accounts are not mutually exclusive, as they may coexist across different brain regions with different neural codes.

Although prioritization has traditionally been viewed as a cognitive computation within WM, recent work has identified active representations of the unprioritized stimulus in the medial temporal lobe (MTL) ^29^, a region traditionally associated with episodic memory but increasingly implicated in WM ^30–34^. Some studies, particularly intracranial recordings in human patients, have also identified other cortical regions, such as the medial prefrontal cortex (mPFC), that contribute to WM ^30, 33^. The MTL and mPFC are part of a broader network, namely the default mode network (DMN) ^35^, whose functional roles in WM prioritization are rarely discussed. Notably, WM-related representations in the DMN are comparably weaker, which may explain why they are most consistently observed using intracranial recordings rather than traditional imaging techniques such as functional MRI (fMRI). On a separate note, the MTL and mPFC are also thought to represent abstract task knowledge in a structured map ^1, 36, 37^, and send this knowledge to the frontoparietal control networks (FPCN) to implement task production. Activity in these regions also tracks task progress and carry information about current goals across multiple trials ^38^. Altogether, these results raise an intriguing possibility that unprioritized task information may be “offloaded” to the DMN for sustained maintenance before returning to a prioritized state, complementary to the FPCN. However, a systematic investigation of this possibility has been lacking.

To summarize, whereas it is widely accepted that prioritized stimulus is supported by a distributed WM network with active neural codes, the maintenance of unprioritized stimulus is more complicated and may involve multiple neural mechanisms. Moreover, despite extensive research on feature-specific stimulus prioritization, whether flexible task prioritization recruits similar mechanisms remains unclear. Recent evidence suggests that task and stimulus information are represented differently in WM ^39, 40^. In particular, the maintenance of abstract task information primarily engages the frontal cortex, whereas maintenance of stimulus information is localized to more posterior sensory regions such as the visual cortex. Maintaining and implementing task information also engage distinct brain regions, with maintenance recruiting more frontal regions and implementation recruiting more posterior regions ^39^. These results suggest that the flexible prioritization among multiple task information may recruit complex collaborations between distinct neural substrates and involve various neural mechanisms. Whether the mechanisms proposed for stimulus prioritization apply to task prioritization requires further examination.

To investigate the neural mechanisms underlying task prioritization during WM, we employed a continuous retro-cue paradigm in which participants switched flexibly between different task rules on a trial-by-trial basis within each block ^41^. Crucially, two randomly selected rules were used per block, requiring participants to retain the unprioritized rule in mind while actively processing the prioritized rule on each trial. Within the trial, participants performed a WM task with two memory delays: the first delay required the maintenance of the task rule and the second delay required the implementation of the rule on a specific stimulus presented between the delays. Combined with fMRI in human participants, this task design allowed us to examine several key questions simultaneously. Specifically, we investigated how prioritized and unprioritized task information are represented during different task stages at the whole-brain level, and what neural mechanisms mediate the flexible switching between different priority states.

## Results

### Task and behavior

Participants (n = 25) performed a delayed manipulation task nested in a continuous retro-cue block design inside an MRI scanner. Within a trial, the task required participants to maintain a rule and a visual stimulus in mind and then to manipulate the latter according to the remembered rule (Figure 1A). Specifically, participants were first cued by a square or a circle to indicate the rule to be used, and maintained the information over the first delay period (Delay 1). After that, they were presented with a to-be-memorized stimulus with a specific size and color (Sample), and maintained both the rule and stimulus over a second delay (Delay 2), before adjusting the on-screen stimulus by pressing buttons during the response period (Response). Furthermore, only two task rules from a set of four possible rules were selected and cued throughout each scanning block (18 trials). At the start of a block, participants were presented with two rules in textual form and two shapes (i.e., square and circle), indicating the relevant rules for the current block as well as the associated visual cues (Figure 1B). After participants reported successful encoding of the rule-cue pairing, the scanning block would begin, during which the cue identity signaled which rule to use for the ongoing trial (prioritized rule), while the other rule was held in memory for future use (unprioritized rule).

**Figure 1.**
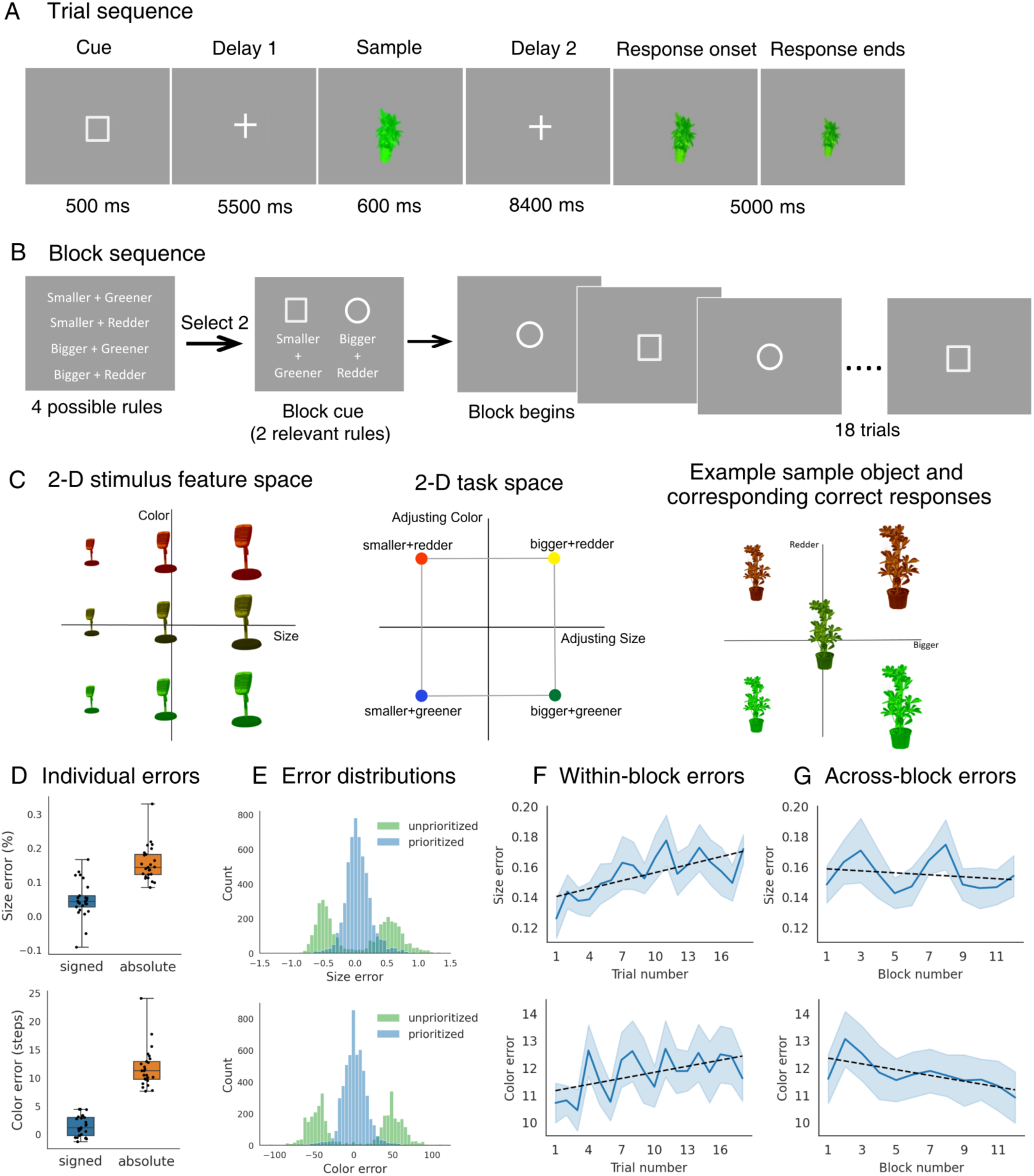
Task schematics and behavioral results. **(A)** Schematics of trial sequence. In each trial, the delayed manipulation task required the memorization of a task rule and stimulus features, and at response phase, the manipulation of stimulus’s size and color based on the cued rule. Response onset was marked by the re-appearance of the stimulus on screen, however, the size and color of this initial item were randomized to prevent motor planning. **(B)** The trial sequence was nested in a block design where participants flexibly switched between two rules across 18 trials. At the block onset, the two selected rules (out of 4 possible ones) along with their associated visual cues were presented on screen. The task would begin after participants indicated successful encoding of the pairing. **(C)** Rule and stimulus designs. Both task rules (middle) and stimuli (left) were constructed from two orthogonal axes. The task space consisted of adjusting size and color to two opposing directions while the stimulus space of continuous stimulus size and color (red-greenness). Right: example sample stimulus (center) and the corresponding correct answers (in four quadrants) according to different task rules. The distances between any given sample and correct answers in terms of feature values were of a fixed value and pre-learned by participants. **(D)** Size (upper panel) and color (lower panel) errors for the fMRI experiments. The response errors were calculated by subtracting correct from responded values. Mean signed error (blue) showed the degree of bias in participants’ responses whereas mean absolute error (orange) showed the degree of precision. Black points represent averaged error of individuals. **(E)** Trial-wise signed error distributions using correct answers (cued rules; blue) and the alternative answers (uncued rules; green). Note that to plot the unprioritized rule distribution one third of trials were excluded due to overlapping dimensions between selected rules (e.g., smaller/greener vs. smaller/redder). **(F)** Memory precision as a function of trial numbers within blocks. Error bands denote standard error of the mean (SEM) using the number of participants as the degree of freedom. Black broken line indicates fitted linear regression trend. **(G)** Memory precision as a function of block numbers.

Stimuli varied along two orthogonal feature dimensions (Figure 1C, left), size (from small to big) and color (from green to red). Correspondingly, task rules (Figure 1C, middle) also varied along two orthogonal rule dimensions, size (adjusting smaller/adjusting bigger) and color (adjusting greener/adjusting redder) changes. Critically, the correct degree of adjustment the participants needed to apply was fixed (Figure 1C, right), such that for a specific rule and a stimulus with particular size and color values, the correct response was definite. Participants learned this adjustment prior to the scanning session. This design allowed for the dissociation between rule and stimulus information, meaning that for different stimuli, the same rule would lead to different correct responses.

Response errors were calculated by subtracting correct answers from responses. Mean signed size error across individuals (Figure 1D) was 4.85% (of starting size, SD = 5.50%), and mean signed color error was 1.50 color steps (SD = 1.82). Mean absolute size error was 15.53% (SD = 5.23%), while absolute color error was 11.80 steps (SD = 3.55), both comparable to our previous studies using a similar paradigm ^39^. These results suggested that participants were able to memorize the correct stimuli and rules and to perform the task according to the instructions. As shown in Figure 1E, distributions of responses error were centered around zero if compared to the correct answers (consistent with the prioritized rules), but formed a binomial distribution away from zero if compared to the uncued rules, confirming that participants made responses following the cued rules. Response error gradually increased across trials within a block (size: *t*(24) = 2.90, *p* = 0.0039; color: *t*(24) = 1.80, *p* = 0.072; Figure 1F), but remained relatively stable across blocks (size: *t*(24) = −0.56, *p* = 0.57; color: *t*(24) = −1.43, *p* = 0.15; Figure 1G).

### Spatial separation of task rule representations in distinct priority states

To investigate the neural underpinnings of task prioritization, we first set out to examine where in the brain multiple rules with different priority states were maintained in WM, using whole-brain support vector machine (SVM) -based multivariate pattern classification for prioritized and unprioritized rules separately. Significant decoding accuracy above chance level would indicate active neural representations that can be read out from BOLD activity in fMRI. The accuracy-based decoding analysis revealed vastly distinct pictures: for the prioritized rule, distributed brain regions across occipital, parietal, and frontal cortices represented the relevant information during both delay periods, consistent with previous work on stimulus prioritization. Furthermore, Delay 1 was characterized by more frontal involvement, whereas Delay 2 engaged more concentrated posterior regions in parietal and occipital lobes (Figure 2A and Supplementary Table 1). In contrast, unprioritized rules were only decodable from posterior visual-related areas in the occipital and temporal cortices, with minimal overlap with those showing prioritized rule representations. This was still true during Delay 2 when prioritized rule representations aggregated towards the posterior cortex (Figure 2B and Supplementary Table 2).

**Figure 2.**
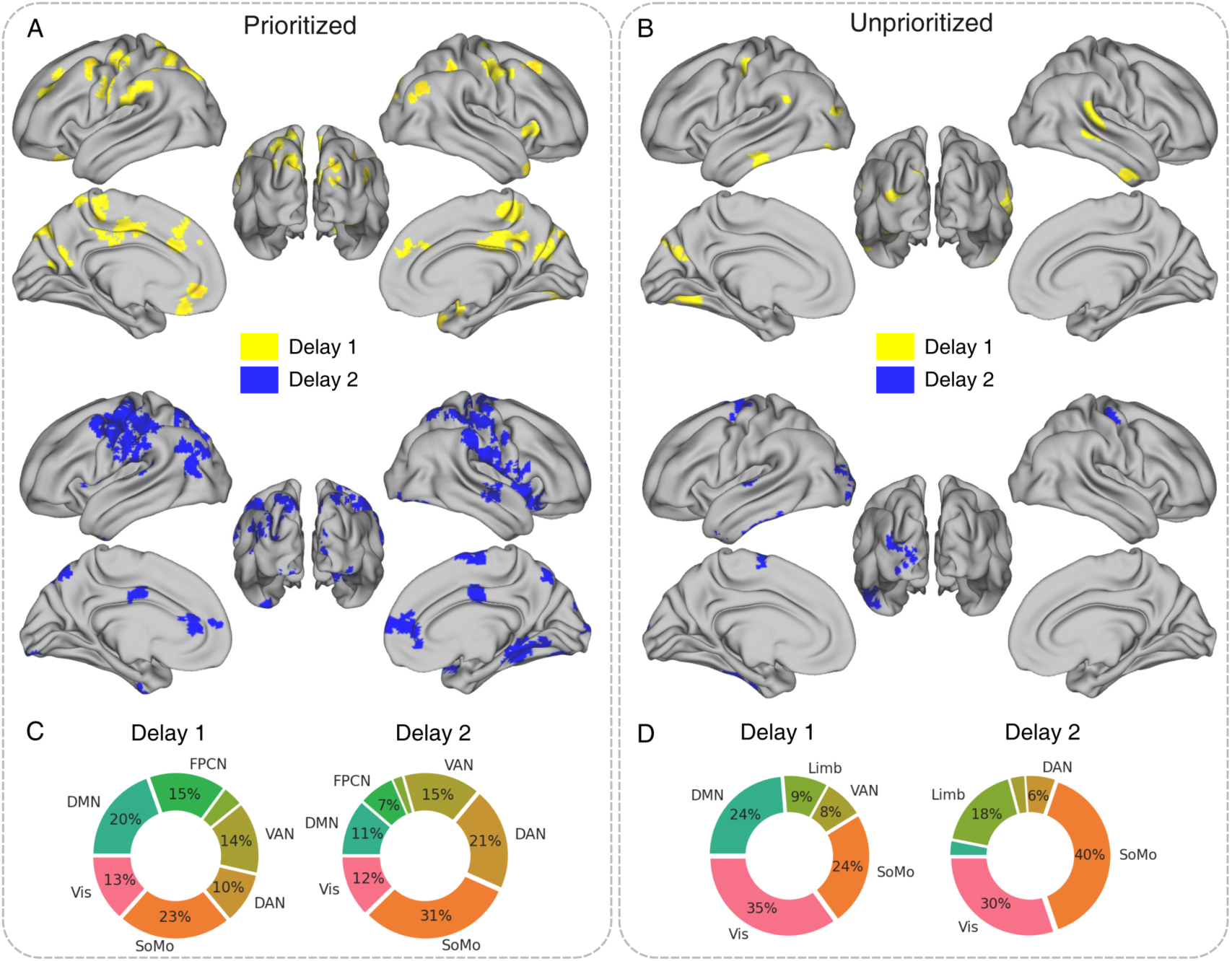
Active storage of prioritized and unprioritized task rules. (A) Cortical regions showing significant above-chance classification accuracy in Delay 1 (yellow) and Delay 2 (blue). A cluster threshold of 80 voxels was applied to the statistical maps for visualization. (B) Same as (A) but for unprioritized rules. (C) Summary of significant regions based on the Yeo 7-network atlas for prioritized rules. Only networks accounting for higher than 5% of all voxels were labeled. (D) Same as (C) but for unprioritized rules. DMN = Default mode network; FPCN = frontoparietal control network; Vis = Visual network; DAN = Dorsal attention network; VAN = ventral attention network; SoMo – Somatomotor network; Limb = Limbic network.

To better compare the results of whole-brain decoding, we summarized the clusters based on the Yeo 7-network atlas ^42^. Figure 2C & 2D showed the proportion of voxels belonging to each key network relative to all significant voxels for a given priority state and time period. As observed earlier, for the prioritized rules, the proportion of voxels was relatively evenly distributed across different networks. Conversely, unprioritized rules were stored significantly more in the visual network compared to the prioritized rules at the same time periods (two-sample proportion test: Delay 1: *z* = 13.67, *p* < 0.001; Delay 2: *z* = 11.39, *p* < 0.001) and no clusters were found in FPCN. Collectively, the decoding results suggested an interesting pattern of spatial separation for the concurrent storage of multiple rule representations: the rule in current use was maintained in a widespread manner, whereby information of prospective relevance was confined to the posterior cortex.

To further understand the nature of rule representations, we analyzed an additional dataset collected from the same individuals in a separate scanning session (all but four participants completed this extra session; n = 21), during which one task rule was signaled by texts (instead of shape cues) from all four available options with equal likelihood at any given trial. This trial-wise design posed no need for the participants to constantly keep another rule or the rule-cue associations in mind while completing the current objective, making it an ideal baseline condition for the primary task (Supplementary Figure 1A & 1B). We identified a similar pattern of single rule representation to that of the prioritized rule in the 2-rule condition, with the single rule maintained in a highly distributed manner in both delays (Supplementary Figure 1C). This indicated that the maintenance of prioritized rules recruited regions similar to those engaged in the 1-rule condition, unlike the unprioritized rules. To examine whether prioritized rules shared neural representational patterns with single rules within these overlapping regions, we trained a classifier using data from the 1-rule task to predict prioritized rules from the 2-rule condition. Successful generalization would provide strong evidence for consistent neural representations across tasks. We found support for this claim in most areas revealed by the separate decoders, reaffirming the correspondence between prioritized and single rule representations. In sharp contrast, repeating the analysis for unprioritized rules yielded no significant clusters (Figure 3A). This demonstrated that unprioritized rules not only recruited spatially distinct regions, but also relied on neural codes that shared no discernable similarity with the single rule. Altogether, these observations suggested that when multiple items were held in WM, prioritized task rules were encoded in a way akin to a condition where only one rule was remembered, whilst unprioritized rules were preserved in a non-overlapping manner.

**Figure 3.**
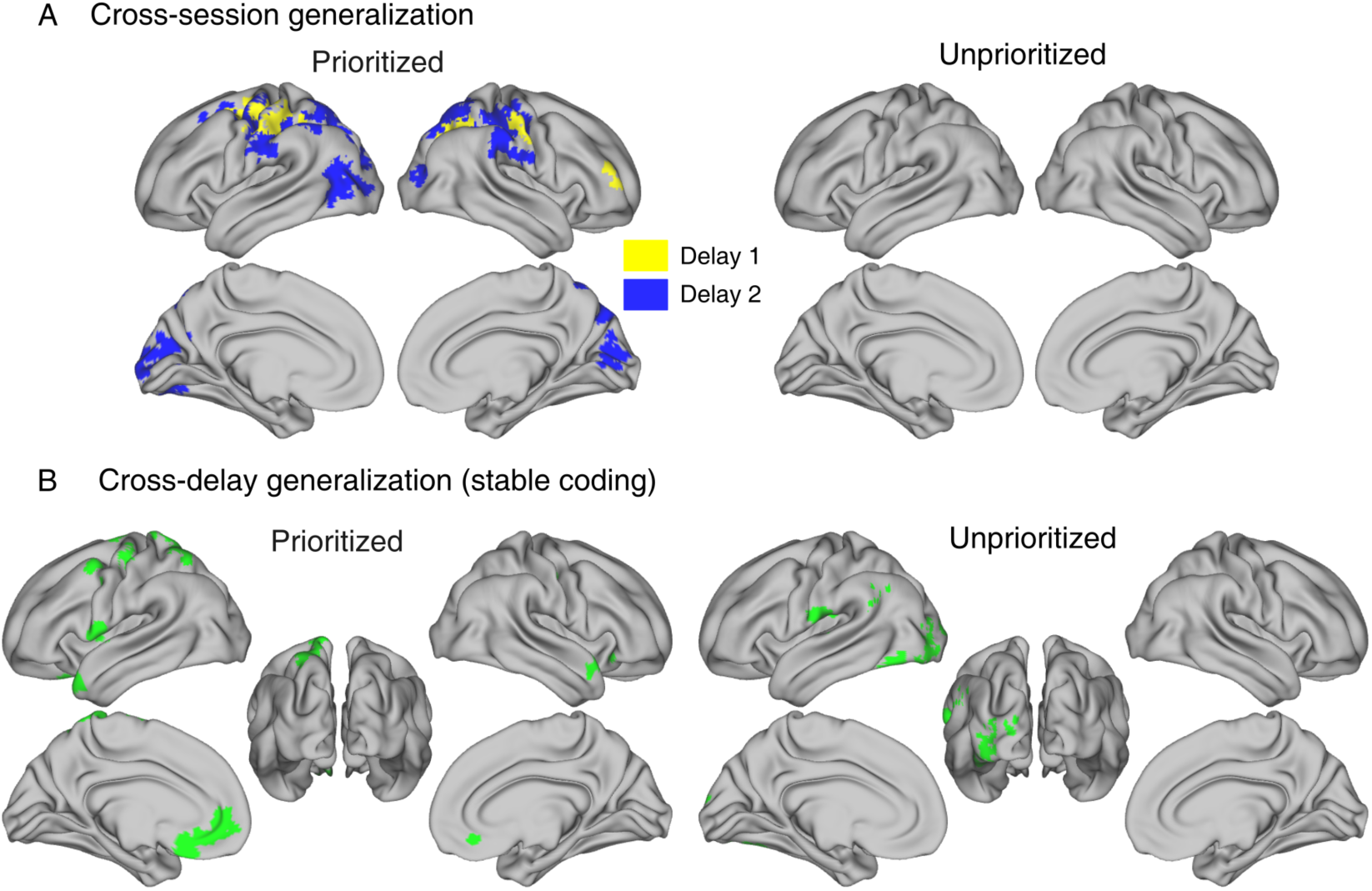
Cross-session and cross-delay generalization. (A) Regions showing generalization between a baseline single-rule condition and the prioritized or unprioritized rules, respectively. Results were intersected by significant self-decoding results in the single-rule condition (Supplementary Figure 1C). (B) Regions showing temporal generalization between two delay periods.

### Spatially separated active representations of prioritized and unprioritized rules support stability within trials

Why does the brain actively maintain prioritized and unprioritized rules in spatially segregated regions? One possibility is that this arrangement was interference-resistant and therefore advantageous to the stable maintenance of both rules. To test this hypothesis, we performed a cross-delay generalization analysis using a distance-based metric to assess whether representations in each priority state were generalizable across the two delay periods. The metric was chosen to mitigate potential influences resulted from rule-irrelevant differences between delays (see Methods). As predicted, subregions within the prioritized and unprioritized networks separately held stable representations of the corresponding rules. Specifically, prioritized rules were stable across delays primarily in more anterior regions, particularly the mPFC, but also including premotor cortex and superior parietal lobe; while stable neural code for unprioritized rules were found in posterior regions, including the left early visual cortex (EVC) and angular gyrus (AG) (Figure 3B). These findings supported the hypothesis that the spatial separation scheme served to maintain multiple rules robustly and minimize potential interference between them.

### Deprioritizing a task relies on a medial DMN-centered network of sustained latent representations

So far, we have observed that memories of multiple rules were mainly retained in a spatially dissociable fashion. An important follow-up question is what mechanisms mediate the dynamic transformation between states. Specifically, when a previously prioritized rule was no longer required, how was its representation transformed to a temporarily unprioritized state (the deprioritization process); and conversely, how did the previously unprioritized item become prioritized when the requirement arose (the prioritization process)? Moreover, would both processes engage the same or separate networks? To address these questions, we tracked the same rule across two neighboring trials when its priority state either stayed the same or changed, and tested where in the brain the rule representation in the first trial would be carried over to the following trial, depending on the type of trials, i.e., whether the second trial was a switch or nonswitch. To better capture potentially subtle covariations in the strength of representations across trials, we chose to use a continuous measure of classifier performance that leveraged its “confidence” in discriminating between trial labels, instead of using binary measures such as prediction success (see Methods). Similar approaches, which afford higher reliability ^43, 44^, have been applied to assess the covariation of neural representations between brain regions ^45, 46^. Specifically, we hypothesized that if a rule underwent a state transition, decoding strength in areas underlying the switching process should uniquely covary with decoding strength in regions where the information was originally represented (Figure 4A).

**Figure 4.**
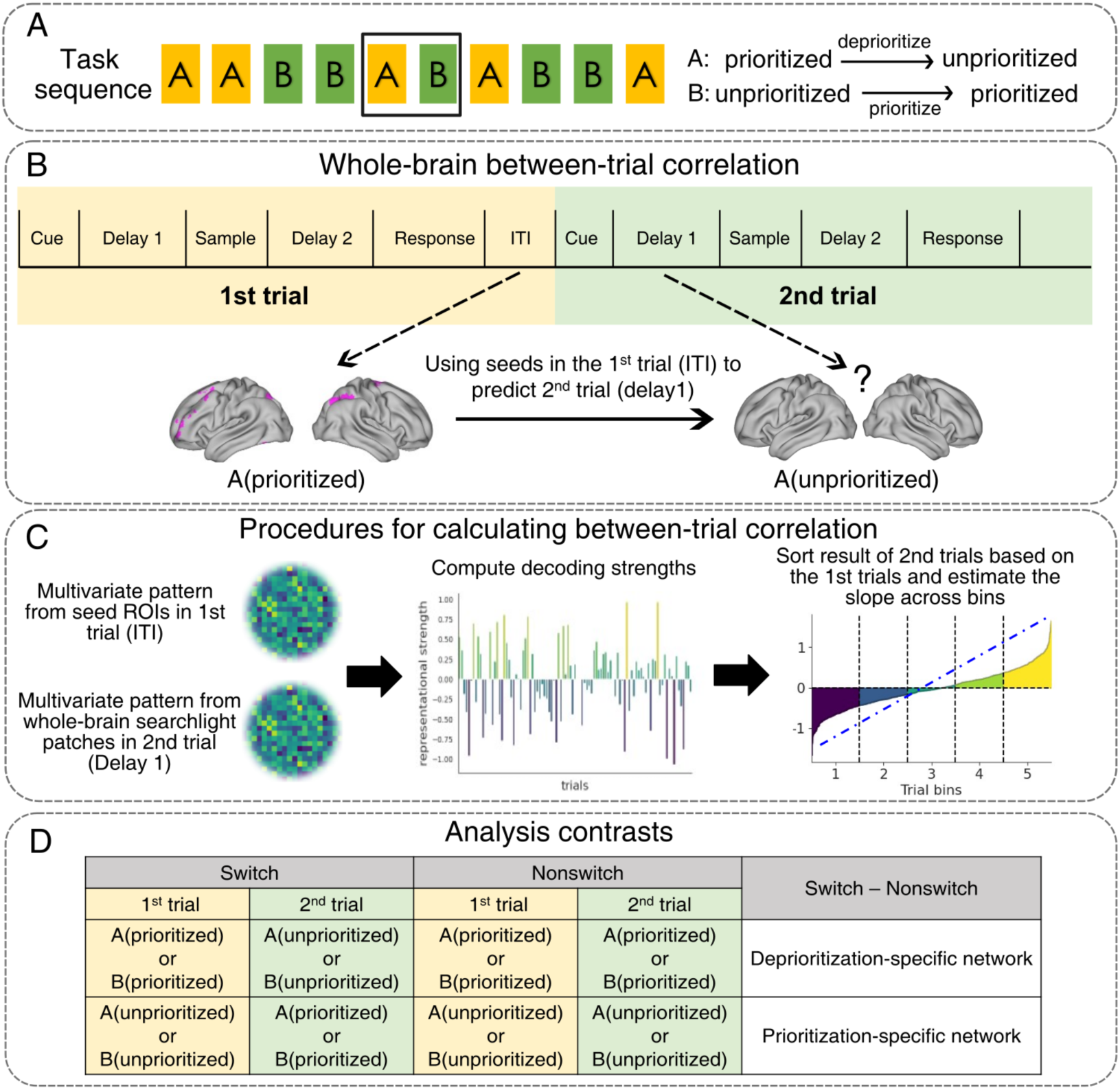
Analysis rationale and pipeline for the flexible switching of priority state. (A) Example task rule sequence and transition of priority state across adjacent trials. Over the course of the two trials highlighted in the black box, rule A was deprioritized and rule B was prioritized. (B) Following the example, to unveil deprioritization-related regions, we first found active representation of rule A (prioritized) during ITI, then used the seed ROIs to predict where in the brain decoding strength of prioritized rule A in the 1^st^ trial was predictive of that of unprioritized rule A in the second trial (switch condition only). (C) Specific procedures for between-trial correlations involved firstly calculating both decoding strengths of adjacent trials, and sorting values of the 2^nd^ trials according to those of the corresponding 1^st^ trials. The rearranged values were divided into 5 equal-sized bins across which the degree of linear trend was assessed. Where in the brain significant between-trial correlations exists, it should be reflected in a positive linear relationship across the trial bins. (D) Rationale of analysis contrasts. To derive networks specific to the switching process, between-trial correlations of nonswitch condition were subtracted from those of switch condition to account for regional connections non-related to priority state transitions.

To this end, we first identified areas where prioritized and unprioritized rules were separately decodable during the inter-trial interval (ITI). ITI was used to capture pre-trial variabilities in rule representations, as activity during this period should reflect a baseline level unaffected by implementation-specific signals (as response period was over) and during which the fate of the formerly prioritized and unprioritized rules were yet undecided. That is, the rule for the next trial could either be the same (nonswitch) or changed to the other (switch), therefore both should exist in a state that was subject to flexible alteration of priorities at this pre-trial phase. Using these ITI clusters as seed regions, we inspected variability in the decoding strength of a task rule as a function of its representation in the preceding trial as well as of state transition, to obtain whole-brain maps of co-fluctuating decoding strength (using beta estimates of Delay 1; Figure 4B). For each seed region, the same rule’s decoding strength in subsequent trials were sorted into five bins according to the ITI decoding strength within that seed (i.e., from lowest to highest), excluding the first trial of each block, and the degree of a positive trend across bins was assessed (Figure 4C). This whole-brain procedure was separately performed for switch and nonswitch trials before contrasting them to obtain switch-specific regions of interest (ROIs; Figure 4D). In a nutshell, the final results revealed brain networks whereby variability in rule representation was uniquely modulated by that of the same item in the preceding trials during a state transition.

During ITI, we found significant representations of prioritized rules (Supplementary Figure 2A) in bilateral fusiform gyrus (FFG), bilateral intraparietal sulcus (IPS), bilateral medial temporal lobe (MTL) including the hippocampus, left dorsolateral PFC (dlPFC) and dorsal middle frontal gyrus (dMFG). Using these seeds, significant covariation in decoding strength between adjacent switch trials were identified in different networks, but most consistently in medial regions including dorso- and ventro-medial PFC (dmPFC and vmPFC), retrosplenial cortex (RSC), and MTL, all key subregions of the canonical DMN ^35^ (Figure 5A and Supplementary Table 3). Post-hoc examinations confirmed that this covariation was only evident in switch but not nonswitch trials (Figure 5B). In other words, the strength of unprioritized rules following a switch was modulated by the same item’s pre-trial representation when it was in a prioritized state. Indeed, DMN showed the highest degree of overlap with results from different seed regions, ranging from 35% for dMFG seed to 98% for IPS seed (Figure 5C and Supplementary Figure 2B). These suggested that medial DMN was particularly involved in deprioritizing a task rule, when the previously prioritized item was no longer required and needed to be offloaded for future use (i.e., a switching-off process), in line with recent proposals on the complementary roles of DMN to the FPCN ^1^. Interestingly, these regions shared little overlap with ROIs showing active decoding of unprioritized rules (i.e., accuracy-based decoding; Figure 2B and Supplementary Figure 3), suggesting that after switching off, unprioritized rules were stored in a distinct set of neural substrates in comparison to the active, on-task storage of the same items. The fact that these regions were not identified by classification accuracy indicated that rule representations in these regions were in a weaker, latent format that was not detectable at the active decoding level, but was distinguishable by measure of decoding strength covariation.

**Figure 5.**
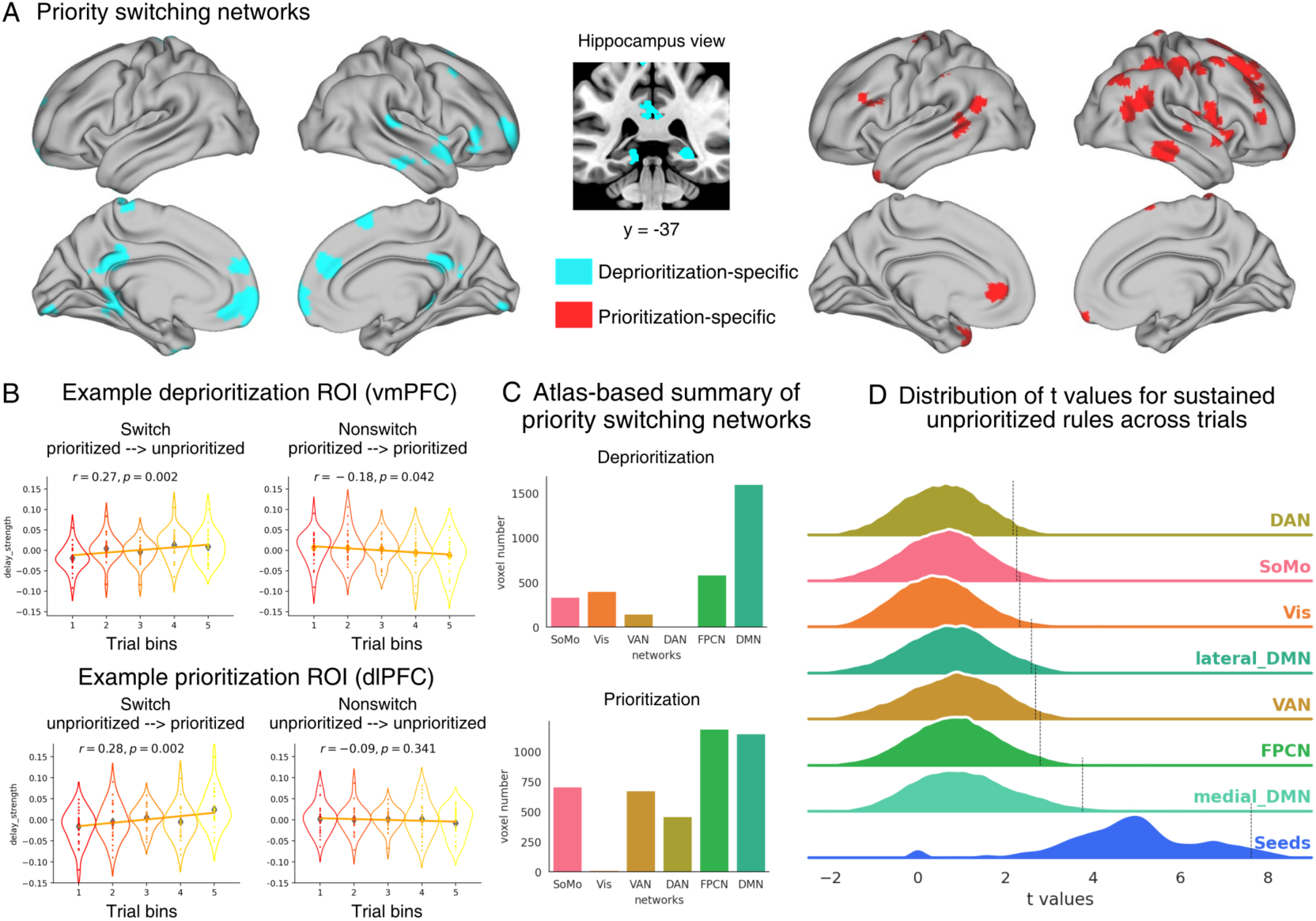
Deprioritization- and prioritization-specific networks. (A) Left: regions showing unique between-trial correlation when a rule was switched from prioritized to unprioritized state (deprioritization; cyan); Right: same as left panel but regions unique to prioritization (red). (B) Upper: distributions of sorted decoding strengths in each trial bins and fitted regression line, separately for switch and nonswitch conditions in an example deprioritization ROI (vmPFC: MNI coordinate = [3, 65, 1]). Lower: same as upper panel but for an example prioritization ROI (dlPFC: MNI coordinate = [29, 34, 39]). Each colored dot denotes one participant’s mean value within bins; diamond markers denote average values across participants. (C) Atlas-based summary of deprioritization- and prioritization-specific networks. (D) The distribution of all group-level *t* values within significant clusters of sustained maintenance of unprioritized information, separated by networks. Black vertical line represents 95th percentile of the sample distribution. Seed: regions that overlapped with the medial-DMN seeds used for this analysis. DMN = Default mode network; Medial DMN: defined as the section of DMN whose MNI x coordinates are within −15 and 15, excluding the seed regions. Lateral DMN: DMN outside of the range for the medial section. FPCN = frontoparietal control network; Vis = Visual network; DAN = Dorsal attention network; VAN = ventral attention network; SoMo – Somatomotor network.

Following this observation, we wondered what functions the latent unprioritized representations served within the deprioritization network. Given that active representation of unprioritized rules supported on-task stability in the visual cortex, it is possible that a second system with weaker representational traces was advantageous to the prolonged maintenance of the temporarily unused information. We tested this assumption using an approach similar to the priority switching analysis, that is, by examining whether there were between-trial correlations of the unprioritized representations across two nonswitch trials, focusing on regions of the medial-DMN within the deprioritization network. Supplementary Figure 4 displayed areas in which unprioritized information was sustained across trials in nonswitch condition, contrasted with switch trials. A few observations could be made: firstly, all seed regions predominately maintained latent unprioritized representations across trials within themselves (rank-sum test, seed vs. medial-DMN: *U* = 7.75×10^7^, *p* <0.001; Figure 5D; for separate seeded maps see Supplementary Figure 4). Moreover, seed ROIs also exhibited a preference in correlating with other medial DMN areas compared to the rest of networks in the parcellation, as well as the lateral section of DMN (*p*s < 0.001), indicating that the deprioritization system was indeed responsible for sustaining latent representations of unprioritized rules over trials.

### Prioritization of task rules engages a different network compared to deprioritization

Using a similar approach, we also investigated the prioritization process. Decoding strength of unprioritized rules during the first trials modulated that of the same item after it subsequently became prioritized (analogous to a switching-on process) in more lateral-focused brain regions such as dlPFC, lateral parietal and temporal lobes (Figure 5A, Supplementary Table 3 and Supplementary Figure 5). Atlas-based network summary indicated that although DMN remained strongly involved, participation from FPCN increased as well (Figure 5C), in accordance with the active decoding results that both networks were involved in maintaining prioritized information. Indeed, unlike the deprioritization process, a high degree of consistency remained between regions actively maintaining prioritized rules and those involved in the prioritization of rules, including the IPS and dlPFC (Supplementary Figure 3). Overall, these findings indicated that the dynamic transition of functional priorities engaged disparate regions depending on the direction of switch: deprioritizing a task rule relied on a latent network primarily consisting of the medial DMN, whereas prioritization-involved areas located predominantly in a lateral network that belonged to a subset of the active representation of prioritized rules.

### Behavioral correlations support dissociable functions of different systems

In the previous section, we identified a latent network centering on the medial DMN that was specifically involved in deprioritization, distinct from the active storage of unprioritized rules. Further analyses demonstrated that this system carried representations of newly unprioritized rules across multiple trials, suggesting a mechanism for sustained, latent storage of temporarily uncued task sets. Functionally, in the present experiment, retaining temporarily unprioritized information throughout a block was as critical as recalling the cued rule for the trial, since the former could be required at any upcoming trial. We thereby reasoned that the representational strength of unprioritized information within the latent network should predict task performance on a longer timescale (e.g., across an entire block), instead of the trial-by-trial processing of active maintenance and implementation. In other words, the overall quality of the unprioritized representations over the course of a block should positively scale with behavior.

To test this, we first fitted a linear model at the individual level using block-averaged decoding strength of unprioritized rules as regressors to predict trial-wise response errors (averaged error across the color and size dimensions; see Figure 6A and Methods), which revealed that in left MTL and vmPFC, variability in block-averaged unprioritized representations was predictive of memory precision (Figure 6B lower). This finding reaffirmed that DMN, especially the medial section, was particularly important to retaining task-relevant information of unprioritized rules in a latent format over longer timescales.

**Figure 6.**
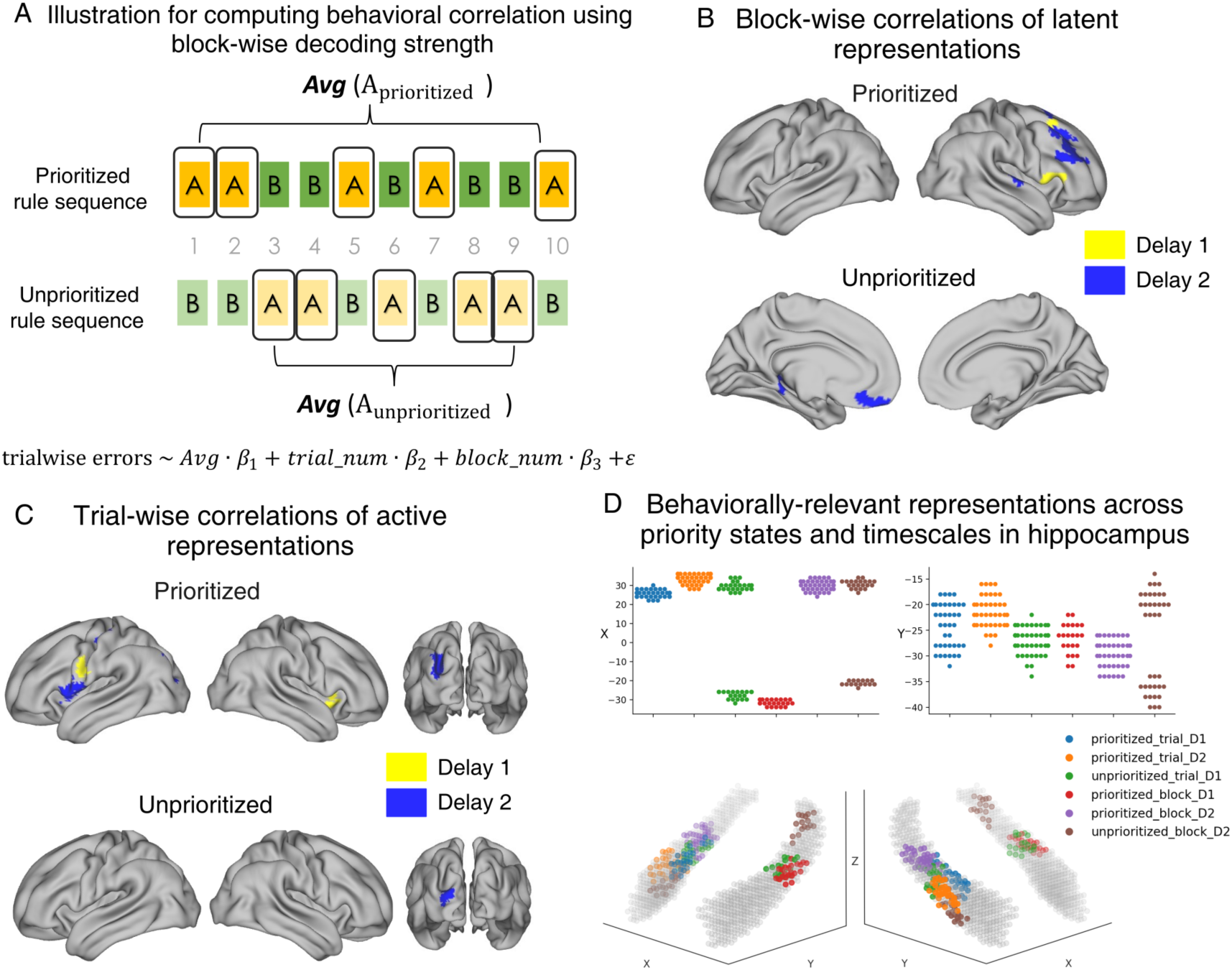
Behaviorally relevant representations of prioritized and unprioritized rules on trial-wise and block-wise timescales. (A) Schematics for computing correlation between trial-by-trial response errors and block-averaged representations. For the example sequence, to test effect of block-averaged unprioritized rules, for trials [1,2,5,7,10], we took *Avg*(*A_unprioritized_*) and for trials [3,4,6,8,9] *Avg*(*B_unprioritized_*) as the predictor values to predict trial-wise response errors. (B) Regions showing positive behavioral correlation of block-wise decoding strengths of prioritized and unprioritized rules. Only clusters intersecting with the prioritization and deprioritization networks (Figure 5A) were displayed, respectively. (C) Same as (B) but for trial-wise representations. Only clusters intersecting with the active representation of rules (Figure 2A & 2B) were displayed, respectively. (D) Behaviorally relevant task representations within MTL. Upper: MNI x and y coordinates of each cluster. Lower: Projections of significant voxels on hippocampus structures. Trial = trial-wise correlations; Block = block-wise correlations; D1 = Delay 1; D2 = Delay 2.

We also probed whether the prioritization-specific network shared a similar function for the prioritized rules. Indeed, within the lateral PFC (lPFC), including the MFG, inferior (IFG) and superior frontal gyrus (SFG), block-averaged prioritized rules positively co-varied with behavior (Figure 6B upper), suggesting that in contrast to the unprioritized counterpart, lPFC was critical for the sustained storage of prioritized information. Taken together, these results demonstrated that the overall quality of both prioritized *and* unprioritized rule representations facilitated performance, albeit through different networks.

Lastly, we assessed whether on-task, active storage of prioritized and unprioritized rules (i.e., accuracy-based decoding) was also associated with participants’ performance on a trial-by-trial basis. Similar to above, we conducted regression analyses but used trial-wise decoding strength as predictors. For prioritized rules, this revealed a number of regions positively correlated with memory precision (Figure 6C upper): right middle insula, postcentral gyrus and IPS. For the unprioritized rules, EVC, which held temporally stable unprioritized representations (Figure 6C lower), was positively associated with memory precision. The finding was consistent with the notion that spatially segregated, active representation of unprioritized rules in the visual cortex was facilitative to the task.

### Hippocampus as a key node for balancing cognitive stability and flexibility

As a key node in DMN, hippocampus has been shown to encode both prioritized and unprioritized items ^29^. In the current study, it was repeatedly indicated in earlier results (Figures 5A & 6B; Supplementary Figure 2A), suggesting a possible multi-faceted role in supporting the maintenance and flexible shift of multiple task rules. We next performed an exploratory analysis focusing on this anatomical region (defined by a subcortical atlas ^47^) to elucidate hippocampus functions in representing rule-specific information, regardless of network constraints. Using a smaller searchlight patch on unsmoothed voxel patterns to avoid contamination of activity from nearby structures, we conducted both trial-wise and block-averaged behavioral analyses within the hippocampus, which revealed a strong involvement across functional states and timescales (Figure 6D). Specifically, higher memory precision was associated with stronger trial-wise prioritized and unprioritized representations. On a longer timescale, memory performance was also positively correlated with block-averaged representations in both priority states. There was no significant difference between prioritized and unprioritized representations in terms of where they situated along the hippocampus long axis (rank-sum test using MNI y coordinates (anterior to posterior axis) of all voxels within the significant clusters: *M*_prioritized_ = −25.32, *M*_unprioritized_ = - 26.83, *U* = 6891.0, *p* = 0.14). However, there was a small distinction in the X coordinates (*M*_prioritized_ = 20.18, *M*_unprioritized_ = 7.15, *U* = 7381.0, *p* < 0.05), as unprioritized rule clusters were found in bilateral hippocampus, whereas prioritized ones were more right lateralized. Overall, activity patterns within overlapping sets of voxels contributed to behavior-relevant representations in both priority states as well as on both timescales, highlighting hippocampus as a key node that not only maintained multiple task rules concurrently but also unified their on-task and sustained storages, in support of the extended nature of the task.

## Discussion

In this study, we investigated how the brain concurrently represents multiple task information and enables flexible switching between them. Our results revealed two key neural organizing principles in this process. First, active maintenance of prioritized and unprioritized rules was spatially separated: prioritized rules engaged a distributed network across cortex, whereas unprioritized rules were confined to posterior cortex. Second, beyond this active coding scheme, recently deprioritized representations were offloaded to a network centering on the medial DMN, which supported the sustained maintenance of unprioritized rules across trials in a latent manner. Behavioral predictions using trial-wise and block-wise representations further support this neural dissociation. Together, our findings unveil a dual neural system with distinct coding schemes that jointly enable cognitive stability during task implementation and flexibility for task prioritization (see Figure 7 for summarized results).

**Figure 7.**
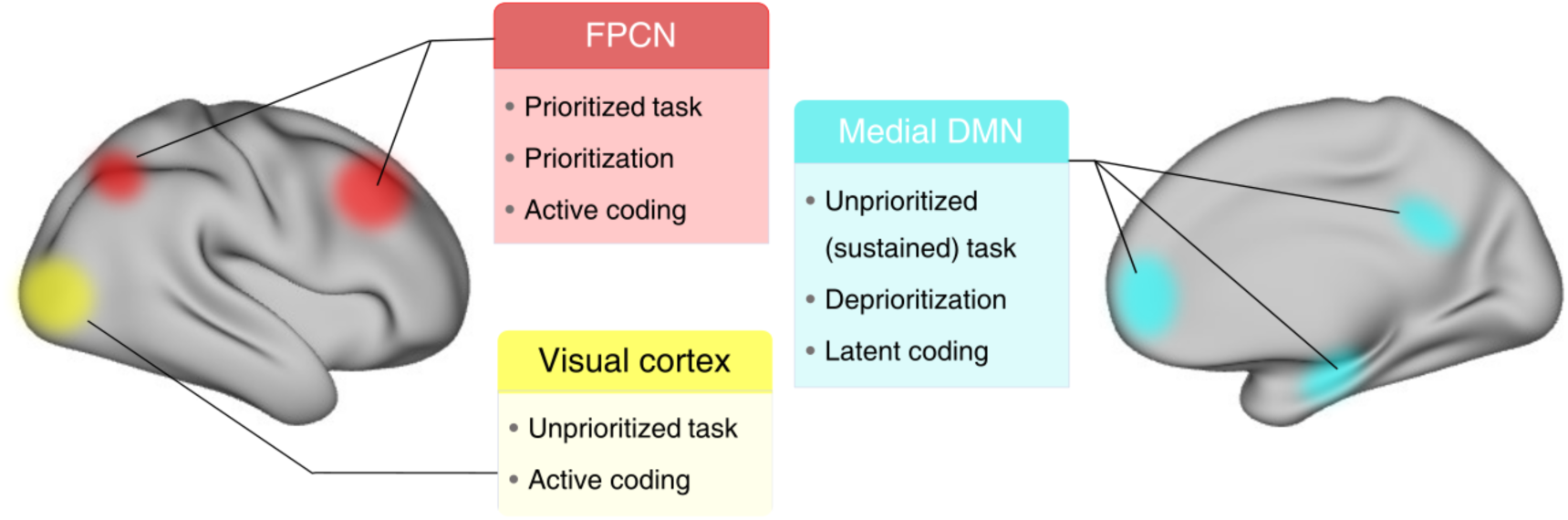
Schematic summary of the main results of the current study. Summary of the key regions highlighted in the analyses for representations of unprioritized and unprioritized task rules, in both active and latent neural codes.

### An active, spatially-separated coding scheme for on-task maintenance of prioritized and unprioritized information

Stimulus-specific delay activity is considered to reflect active representation of WM ^15–19^. With this logic, previous research has identified a distributed cortical network of remembered and prioritized stimulus information during WM ^14–16^. Using a similar multivariate approach, here we demonstrated that prioritized task rules were distributed across the cortex, encompassing sensory, parietal, and frontal cortices, suggesting that active distributed representations serve as a general mechanism for WM prioritization, regardless of the specific types of information being maintained. Reversely, representations of unprioritized rules were less pronounced and restricted to posterior cortex, particularly the visual areas. This account was further supported by the stability of neural codes across delays: prioritized rules maintained stable representations in the frontal and parietal lobes, while unprioritized rules were stable in the visual cortex. Intriguingly, previous work has shown that the frontoparietal cortex, but not the visual cortex, represents unprioritized stimulus information ^13^. In other words, unprioritized representations rely on distinct neural substrates, with the frontoparietal cortex supporting unprioritized stimuli and visual cortex supporting unprioritized tasks. Together, these results point to a unified principle of spatially-segregated active representations for concurrently held WM contents, irrespective of their categories (i.e., abstract task variables or sensory-based stimuli), although the specific neural loci of such schemes may vary depending on the content.

Additionally, our results showed that the spatial distribution of task representations varied with the task stage. When the cued rule was being implemented during the second delay period, prioritized representations shifted more posteriorly. This finding aligns with our previous report that task representations moved towards the visual cortex during task implementation, in order to integrate with stimulus information stored in the same regions ^39^. Combined, these results suggest that prioritized task information is encoded by active, distributed representations, whereas unprioritized task information is represented by separable, localized brain regions.

### A latent coding scheme for sustained maintenance of unprioritized information across trials

Apart from the spatial separation principle in support of representational stability within trials, we uncovered a second coding scheme which utilizes latent neural codes undetectable by standard decoding methods. Recent advance in human single-neuron research has shown that unprioritized stimuli are encoded by persistent neuronal activity in the MTL, although it is comparably weaker than the prioritized stimuli ^29^. This suggests an alternative possibility that unprioritized information may be encoded in weak neural codes that fail to reach significance in standard decoding methods. By developing a more sensitive approach to probe decoding-strength-based covariations, our results reveal a medial-DMN-focused network involved in the switching-off process during task switching, where a previously prioritized rule transitions into an unprioritized state. The identification of DMN instead of FPCN may be surprising at first glance, since they have often been pitted against each other as the “task negative” and “task positive” networks, respectively. However, our results align well with the evolving view on the role of DMN in higher-order cognition, in particular those guided by information from memory ^48^, such as decision-making processes guided by preceding trials ^49, 50^ or task schema ^51^ (), and those involving cognitive transitions, including event boundaries during natural stimuli processing ^52^ and cued task switching between hierarchical nested task sets ^53–55^. Together, our results indicate that DMN engagement could be a hallmark for generic transitions between task demands, although more specific conditions have been discussed to elicit DMN involvement^56^.

Why is unprioritized task information maintained in two distinct systems of neural substrates, each realized by a unique coding scheme? Although we found evidence for relevant information in both the posterior cortex and a more expansive medial-DMN-centered network, the former is a stronger type of representational trace that exceeds decoding threshold, while the latter relies on finer-grained variations in decoding strength. This difference in intrinsic signal strength is potentially related to the divergent functions the systems support. The latent information within medial DMN was found to carry over across multiple trials when the unprioritized rules remain uncued, most strongly within the regions themselves, followed by other nodes of the medial DMN. This highlights it as a relatively self-contained system that sustains unprioritized rules on an extended timescale. The muted nature of this representation may also be energy-conserving over longer time periods and minimize interference with prioritized rules. This claim is further supported by the brain-behavior correlation that regions within the medial DMN predicted memory performance at a block-wise level, corroborating that its contribution is unique to the prolonged maintenance of information rather than on-task processing within each trial ^41^. Perhaps relatedly, subregions of mPFC predicted future strategy before it was actually realized by participants ^57^, raising an intriguing possibility that representations in the mPFC may not necessarily require conscious access, thereby providing a basis for latent, subthreshold format of unprioritized information. Conversely, the strength of active unprioritized representation in the visual cortex scaled with behavior on a trial-by-trial basis, implying it is more linked to on-task computations, possibly reflecting the degree of spatial separation from prioritized items. Moreover, the “redundancy” of retaining the same information through two distinct coding schemes could potentially alleviate memory decay associated with deprioritization, such as those observed for the unprioritized items in a dual retro-cue task ^20, 26, 29^.

### Flexible switching between priority states recruits medial- and lateral-focused systems

In contrast to the deprioritization-specific network, using a similar between-trial approach to examine the “switching-on” process, we instead observed a major contribution from the lateral-focused FPCN. This network overlapped substantially with the distributed, active representations of prioritized rules, implicating that prioritization and on-task active maintenance of the prioritized contents likely depend on unified neural substrates. In turn, this also lends credence to our claim above that the deprioritization process invoked a secondary system different from the active storage of unprioritized rules in the posterior cortex, as they relied on the same analysis pipeline. Our finding that key regions of the lateral frontal and parietal cortices were engaged during dynamic task prioritization aligns with previous task-switching literatures, which similarly identified a lateral PFC-focused system along with other subcortical sites, likely related to fundamental processes these regions serve, such as maintaining task representations and updating content in WM ^58–60^. Crucially, we offered further insights from an informational perspective by showing that shifting unprioritized rules into task focus relied on representing the relevant rule information within FPCN regions. Notably, lPFC within this network predicted behavioral performance for prioritized rules, confirming its functional relevance. Thus, examining the priority transition process across adjacent trials revealed two distinct, direction-specific networks that together enable flexible switching of task rules.

Collectively, the medial- and lateral-focused systems are reminiscent of the recent theoretical framework unifying the functions of FPCN and an MTL-mPFC-OFC (MTL-OMPFC) network, in representing task information to support behavioral flexibility ^1^. Specifically, MTL–OMPFC stores abstract task knowledge in a relational cognitive map suited for generalization, whereas the FPCN integrates this knowledge with other task-related features to implement production (e.g., action or decision-making). The apparent contrast between the prioritization and deprioritization systems in the present study is in strong agreement with this framework: the lateral-focused system is functionally related to switching a task set to a prioritized state for implementation purpose; in contrast, the medial DMN which encompasses MTL-OMPFC is involved in the prolonged preservation of task representations not in immediate use, complementing the role of FPCN. Our results extend the framework by showing that the collaborative contribution of FPCN and medial DMN could be a universal principle in complex task-driven cognition: whereas both networks jointly support computations, the specific functions they subserve are adaptive to the task demands. On a related note, between-network coordination is made possible by the distributed and juxtaposed nature of their topographical organizations, which enables interactions through heterogenous connectivity between subnetworks of FPCN and DMN ^61^.

### Hippocampus as a hub for integrating information across priority states and timescales

We have also showed in hippocampus an interesting aggregation of behaviorally relevant task representations across priority states and timescales. Intracranial recordings of single-neuron activity and local-field potential from MTL have provided evidence for delay-period stimulus-specific persistent activity that is predictive of, and even causal to, WM fidelity ^30–32^. Yet, due to a lack of whole-brain examination, MTL is not well integrated in the current neural models of WM prioritization, which primarily concerns distributed cortical regions. Our study provides evidence for hippocampus’ association with WM using a coarser level of imaging, which can be partially attributed to our block-wise design that extends the task timeframe beyond what is typically used in WM paradigms (i.e., sub-10 seconds). Most crucially, we demonstrated that hippocampus maintained concurrent behaviorally-relevant information of both prioritized and unprioritized rules in a latent format, consistent with previous single-neuron findings ^29^. In that work, image-selective cells within MTL encoded both attended and unattended items, while further divisions of activity profiles among these neurons separated the priority states. It is possible that a similar neuronal mechanism could account for how overlapping sets of hippocampal populations (voxels) in the present study maintained both task rules simultaneously. Overall, our findings highlight the importance of considering and integrating hippocampus functions into the theories of WM-supported goal-oriented cognition.

### Relationship with alternative accounts of WM prioritization

How do the current findings advance our understanding of WM prioritization? By using task rules instead of sensory-based stimuli, we extended previous findings by showing that multiple coding schemes coexist during WM prioritization. While active representation supports the concurrent maintenance of prioritized and unprioritized rules through spatial separation, a latent coding scheme, primarily involving the medial DMN, supports the sustained maintenance of unprioritized information across trials. This latent scheme is related to but different from the activity-silent account: both can offer explanations for the absence of detectable signals in standard multivariate decoding ^20–22^; however, the former still relies on fluctuations in the signals of relevant information, albeit at a weaker, subthreshold level not distinguishable by traditional binary decoding. It should be noted that our findings do not exclude the activity-silent mechanism, as the current study did not measure memory-related synaptic changes ^25^. Rather, the weak coding scheme may help reconcile inconsistencies in the WM prioritization literature, particularly studies that report an absence of decoding for unprioritized information. Specifically, our results suggest that the detection of unprioritized signals requires more sensitive measures compared to traditional practice, yet it remains achievable using noninvasive methods. This is also in line with previous work in which strengthening the signal-to-noise ratio either by maximizing the number of subjects ^13^ or by focusing on a specific oscillatory band ^62^ has led to a reliable readout of unprioritized items. In summary, our findings offer a potentially more sensitive approach that can successfully detect unprioritized memory content using noninvasive measurements in humans. Finally, this study did not directly probe the recoded representation account, as the two rules were always different on every trial, resulting in an inherent anti-correlation between rules that could drive opposite representations. Future studies using modified task designs ^26, 27^ can help to further address these issues.

### Conclusion

In summary, we identify two concurrently operating neural coding schemes that together support flexible task prioritization and deprioritization, highlighting a fundamental network mechanism that enables human multi-tasking cognitive flexibility.

## Materials and methods

### Participants

Twenty-five MRI-eligible participants were recruited for the experiment (mean age = 24.3 years; age range = 21-30 years; 20 females). Twenty-one of them also participated in an additional scan session on a different day performing an alternative version of the delayed-recall task (mean age = 24.2 years; age range = 21-30 years; 16 females). All participants were recruited from the Shanghai Institutes for Biological Sciences community, reported neurologically and psychologically healthy, had normal or corrected-to-normal vision, provided written informed consent, and were monetarily compensated for their participation. The study was approved by the Ethics Committee of the Center for Excellence in Brain Science and Intelligence Technology, Chinese Academy of Sciences (CEBSIT-2020028) and conducted according to the principles in the Declaration of Helsinki.

### Experimental design and procedures

#### Overview

Participants performed a delayed manipulation task nested in a continuous retro-cue block design. On each trial, the delayed manipulation task required the maintenance of both a task rule and a specific stimulus and at later stage the manipulation of the stimulus features according to the rule. Specifically, each stimulus varied along two dimensions, size (small to big) and color (green to red). Correspondingly, there were four task rules of size and color manipulations: changing the stimulus to be **1)** bigger and redder, **2)** bigger and greener, **3)** smaller and redder, and **4)** smaller and greener. Only two task rules were selected and cued throughout each scanning block (18 trials). Task cues alternated between the two rules across trials randomly, creating two priority states for every trial (prioritized vs. unprioritized).

#### Definition of stimulus features

Stimuli generation and presentation were implemented using PsychoPy (version 2021.2.3) ^63^. Stimulus color (greenness-redness) was generated by first converting the images into grayscale to remove all original hues while preserving the relative opacities of pixels using the Image module from Python package PIL. To make the feature transition from extreme greenness to extreme redness in a smooth fashion, the resulting files were converted to RGBA mode and the green-redness values were manipulated by adding/subtracting the same value in the R and G channels whereas the value in B channel was set to zero. The RGB value range for PsychoPy was 0-1, each button press moved the R and G value of all image pixels to opposite directions by 1%. The values in R and G channels were rescaled to 0.25-0.75 with a mean of 0.5. Therefore, for each stimulus there existed 150 variations of color-manipulated images (until all pixels had a value of 0 or 1). However, since all pixels appearing uniformly green or red would render the stimulus unidentifiable (because there is no contrast), the 30 images with most extreme R/G values were discarded, resulting in 120 usable color-varied images - these made up 120 adjustment steps for the color dimension. Stimulus size was directly controlled in PsychoPy using the size attribute and the allowed range was set to 0.01-0.45 screen height. Both starting size and colors were drawn from three predetermined bins, resulting in nine unique conditions. For size the bins were 0.17±0.01, 0.22±0.01 or 0.27±0.01 of screen height; for color they were 34±2, 58±2 and 82±2, indexing from the 120 color steps explained earlier. The distance between each particular sample stimulus and its corresponding correct answer for size was ±24% of the original size (unit: screen height), and ±26 (unit: color steps) for color. Importantly, the degree of this distance in both features were fixed and independent of the initial stimulus values or the rule, making the responses unrelated to other task variables. Participants learned the required distance during the behavioral learning session.

#### Behavioral learning

In the behavioral session, which was held one or two days before the scan session, participants learned the distance of required adjustment by first viewing all pairs of starting and target values in size and color shown side-by-side, respectively (162 trials per stimulus feature), before receiving a Two-Alternative-Forced-Choice (2-AFC) test whereby they were given a starting stimulus and asked to choose the correct target stimulus from two options (80 test trials per stimulus feature). Next, both features were combined together in the same stimulus in another round of 2-AFC test to familiarize them for simultaneous adjustment of both size and color based on one of the four rules, in preparation for the main task (100 trials). Finally, participants completed 180 trials of the main task in order to apply the learned distance of adjustment to stimulus features. Trial-wise feedback of response error was given in the behavioral session only so that participants could keep improving their performance.

#### Main task

Each trial in the delayed manipulation task consisted of two memory delays and a response period. The first delay required the maintenance of the rule only, and the second delay required both the maintenance of the rule and the stimulus. At the beginning of a trial, a shape cue appeared centrally on the screen for 500-ms (Rule cue), signaling the direction of manipulation in the current trial. This was followed by a 5500-ms delay (Delay 1), during which participants should retain the manipulation rule in mind. Following the first delay, a stimulus object was presented for 600 ms (Sample), which was randomly chosen from exemplars belonging to three conceptual categories with similar outlines: bowling pins, plants and microphones, provided and detailed in ^64^. Participants were instructed to maintain the object’s size and color during a second delay period for 8400 ms (Delay 2). The relatively long durations were set to accommodate the sluggishness in hemodynamic responses. To prevent participants from using physical changes on the retinal image or distance to the edge of monitor as memory aids, the stimulus location in each trial was randomized along an invisible circle around the center of the screen with a radius of 0.08 monitor height (i.e., a 2° visual angle difference from the center). During the response phase (Response), the same object reappeared with a random but different size and color from the remembered values, in order to prevent motor preparation, as participants could not know in advance which directions to move the on-screen stimulus to achieve the correct values. They were given a maximum of 5000 ms to adjust the stimulus features to the correct values using button presses. The correct response depended on both the starting values and the cued rule, as learned in the behavioral session. Thereafter, inter-trial intervals (ITI) were jittered between 4000, 5500 and 7000 ms with equal likelihood.

This delayed manipulation trial structure was nested in a block design, within each block of 18 trials, only two of the four task rules were available to be cued. Specifically, at the beginning of every block, two rules paired with two shape cues (a square and a circle) were shown for participants to memorize (Figure 1B). After participants indicated successful remembering of the associations by verbal confirmation (typically took 5 to 10 seconds), the block would start with the two presented rules alternating between trials at random. The same visual cues were re-used in every block with different cue-rule associations, thus disentangled from task rules. Within blocks, rule types and stimulus feature bins were counterbalanced, while across blocks task rule combinations (i.e., the pair of rules used in a block) were randomized and each repeated twice. Relations between pairs of rules and the visual cues across the two repetitions were also counterbalanced. Overall, participants completed 12 functional blocks lasting 466.5 s each, totaling up to 216 trials.

#### Single-rule task (baseline condition)

In a separate session, participants performed an additional task in which only a single task rule was recruited on every trial with no unprioritized rule involved. The single-rule task shared the same trial structure as the main task, except that the shape cues were replaced with textual cues. At the beginning of each trial, participants were presented with a textual cue indicating the rule to be maintained and implemented in the current trial, with rules counterbalanced across trials. Participants completed 216 trials of the single-rule task divided in 12 runs, same as the main task. Data from the single-rule task have been reported previously using a different analytical approach ^39^. The order of the two tasks were counterbalanced across participants, with half performing the single-rule task in their first scan session and the others in their second.

#### Data acquisition

MRI scanning was performed at the Brain Imaging Center at Institute of Neuroscience, Center for Excellence in Brain Science and Intelligence Technology, Chinese Academy of Sciences, on a Siemens 3T Tim Trio MRI scanner with a 32-channel head coil. High-resolution T1-weighted anatomical images were acquired using a magnetization-prepared rapid gradient-echo (MPRAGE) sequence (2300 ms time of repetition (TR), 3 ms time of echo (TE), 9° flip angle (FA), 256 × 256 matrix, 192 sequential sagittal slices, 1 mm^3^ isotropic voxel size). Whole-brain functional images were acquired using a multiband 2D gradient-echo echo-planar (MB2D GE-EPI) sequence with a multiband acceleration factor of 2, 1500 ms TR, 30 ms TE, 60° FA within a 74 × 74 matrix (46 axial slices, 3 mm^3^ isotropic voxel size). After four experimental runs whole-brain Fieldmap phase and magnitude images were collected for correction of EPI distortions. Stimuli were projected on a 1280 × 1024 resolution MRI-compatible screen at the back of the scanner. Participants viewed the screen through a mirror attached to the head coil with a viewing distance of 90.5 cm, and used two two-button response boxes, one in each hand, to adjust the stimulus features.

### Data preprocessing and analyses

#### Data preprocessing

Preprocessing of MRI data was performed using fMRIPrep 21.0.2 ^65^, which is based on Nipype 1.6.1 ^66^. For each experimental run, the single-band reference data were taken as the reference volume. A B0 nonuniformity fieldmap was estimated and aligned to the reference volume. The reference volume was corrected for distortions using the fieldmap, and was co-registered to the anatomical scan. Both the anatomical and functional scans were then normalized to the MNI152 template. A detailed description of preprocessing procedure generated by fMRIPrep can be found in Supplementary text.

#### Single-trial general linear model (GLM)

A single-trial-level univariate GLM was fit to extract neural response estimates for each delay period while accounting for hemodynamic response delay. For each functional run, the design matrix included task regressors representing separate trial stages (e.g., Rule cue and Sample) and trial-wise regressors associated with the period of interest. For example, for modelling beta values of Delay 1, the matrix columns were comprised of Rule cue (500 ms), Sample (600 ms), Delay 2 (8400 ms), Response (5000 ms), and trial-wise regressors for Delay 1 (i.e., Delay1_trial1[5500 ms], Delay1_trial2, etc). The procedure was repeated for other trial periods of interest. Thus, each trial was separated out into its own regressor of interest within the design matrix ^67^. Additionally, the model included six head-motion regressors, three global signals from CSF, white matter and global activity, and three trend predictors from a polynomial drift model. Task regressors were convolved with the SPM canonical hemodynamic response function (HRF) and its time derivatives. Functional data were standardized, high-pass filtered at 0.01 Hz and spatially-smoothed with a 4-mm FWHM kernel before model fitting. The resulting trial-wise beta series were used for all subsequent analyses.

To estimate neural activity during the ITI, we adopted the method employed by previous work ^38^ and used an additional regressor, i.e., a stick function placed at 4000 ms after response offset, to model the ITI-related betas, which was the last TR before the next trials began for those with the shortest ITIs. All other procedures remained the same as the delay-related single-trial beta estimation.

#### Whole-brain and hippocampus searchlight procedures

For an array of results presented in the current study, we combined various analyses (as detailed below) with a searchlight procedure ^68^ in order to identify regions related to different functions across the whole brain. Using the Searchlight function in Nilearn, a spherical patch with a 9-mm radius centered on each voxel was constructed within the whole-brain gray matter mask. For searchlight within the hippocampus, we instead used a spherical patch with a 6-mm radius. The definition of the hippocampus region of interest (ROI) was based on a subcortical atlas ^47^.

#### Cluster-level permutation-based multiple comparison correction for searchlight results

We carried out group-level cluster-based multiple comparison correction for all searchlight analyses results. We first generated a null distribution of the group mean by running the above searchlight procedure with shuffled trial labels 100 times for each participant. Then the permuted maps were bootstrapped from each participant 10000 times to create a null distribution of group-level mean ^69^. Thirdly, the true group-level statistics (averaged across individual maps) were compared to the null distribution to compute the voxel-wise p-values. All voxels that passed the significant threshold (α = 0.05) and formed clusters, defined as neighboring by face only, were identified. Finally, we determined cluster-level threshold by iteratively drawing group-level shuffled maps from the null distribution to obtain the largest continuous cluster size using the same criteria in Step 3. This was repeated 10000 times to generate a null distribution of cluster statistics, from which the 95th percentile value was used as the cluster size threshold. Furthermore, for all cortical brain results shown in figures we adopted a visualization cluster threshold = 80 voxels, except for the hippocampus-focused analyses, for which we used a liberal threshold of 20 voxels to adjust for the relatively small ROI size, and to avoid biasing the final interpretations. Visualizations were done using tools from Connectome Workbench ^70^.

#### Accuracy-based decoding

We explored where the prioritized and unprioritized rules were maintained by means of multivariate pattern decoding. Using the Scikit-Learn SVC class, decoders were trained on beta values from voxels within searchlight patches based on either the prioritized or unprioritized rule labels (i.e., the underlying data were identical). A special cross-validation process was adopted in which two blocks with non-overlapping rule combinations (e.g. [bigger & redder, bigger & greener] for block 1 and [smaller & redder, smaller & greener] for block 2) were held out as testing set in each fold, as opposed to the traditional leave-one-run-out procedure. This was to ensure that trial numbers of each condition were balanced due to the block design. Results were assessed by accuracy averaged across folds and compared to the 25% baseline (one out of four rules). For cross-session decoding analyses, we trained decoders using all trials from the single-rule session, and tested them on the 2-rule data. Visualization of results was further intersected by significant self-decoding results in the single-rule session to ensure that all reported clusters contained significant single-rule representations (as a prerequisite to shared neural codes).

#### Distance-based decoding

Unlike the straightforward multivariate decoding of rule conditions, the test for stable (cross-delay) neural codes involved evaluating generalizability between two sets of data. This may potentially be problematic for accuracy-based metrics as it can be degraded by irrelevant signal differences between the two patterns even if rule-related signals were perfectly overlapping. For example, if there were additional delay-specific activities which were not fully orthogonal to the axis that differentiated task rules, generalization as measured by accuracy could be hindered below baseline despite of a shared representation across delays. Due to this consideration, we utilized an alternative distance-based metric that has been previously applied to similar generalization testing ^71^ to examine cross-delay rule representations. Specifically, the metric was defined as the difference in trial-averaged distance to the fitted SVM’s decision hyperplane between two conditions. Since there were four task rules, we further averaged the difference across all six possible pairs of conditions as the decodability measure. This method is more suited for generalization testing, as it is less influenced by non-orthogonal signals in the held-out data than accuracy-based decoding, as long as there remain systematic differences in their respective distances to the decision boundary. Reversely, if the training and testing sets are truly non-generalizable, the averaged distance contrast would theoretically be zero. Of note, the final outcome was acquired by averaging over cross-decoding of both directions to avoid biases in the decoders trained on only one set of data, i.e., trained on Delay 1, tested on Delay 2, and vice versa. To ensure consistency across different decoding metrics and to assist interpretation, the ROIs identified from the whole-brain cross-decoding analyses and cluster-level thresholding (Supplemental Figure 6A) were subsequently cross-validated in the following steps: first, we subjected the neural activity patterns of each ROI to a permutation test ^71^ to verify results from whole-brain analyses. Instead of using zero as the theoretical baseline, we generated group null distributions by shuffling the testing set labels before calculating the distance contrast for 5000 times. Clusters with real values below the 95th percentile of the resulting null distribution were excluded from further analysis. This method not only ensured the validity of ROIs by using an experimentally-generated null and threshold, but also bypassed the high computational cost of computing the null distributions for the whole-brain maps. Second, to ascertain that ROIs showing cross-decoding results also exhibited significant self-decoding (i.e., trained and tested on the same delay period), we also repeated the first step with decoders trained and tested on the respective sets of data (cross-delay) alone. For instance, for all clusters with shared representations between delays, we trained two separate decoders with either Delay 1 or Delay 2 beta values using the distance metric to confirm that neural patterns within these regions indeed held information of task rules. Regions that did not meet this requirement were also excluded. In summary, the final ROIs needed to show both significant self- and cross-decoding, assessed by the rigorous permutation-based test (Supplemental Figure 6B and 6C).

#### Priority switching-related analyses (between-trial correlations of decoding strength)

To understand how rule representations dynamically changed as a result of shift in their priority and what neural substrates supported this cognitive flexibility, we tracked the fate of the same task rule across two neighboring trials, separated by whether a switch occurred in between (switch vs. nonswitch). Importantly, in keeping with previous studies that reported higher reliability with continuous measures of classifier performance ^43, 44^, we used a more refined variant termed “decoding strength” that leveraged the decoder’s “confidence” in discriminating between trial labels instead of using binary measures such as prediction success (0 or 1). Specifically, we first estimated the probabilities of the decoder in predicting labels of the held-out trials by setting the parameter *probability* = True in Scikit-Learn’s SVC function. This returned one value for each condition label per trial, from which we calculated the difference between the probability associated with the correct label and the highest incorrect label. To note, this metric would be negative if the predicted label was incorrect. Given that a decoder can classify two trials accurately albeit with different levels of confidence, this method allowed a finer-grained quantification of representational strength by taking into account the discriminability of trials and have been applied in similar analyses assessing the covariation of information between brain regions ^45, 46^. The continuous nature of the decoding strength measure also naturally lend itself to the present analysis which assessed the co-fluctuation of representations between two adjacent trials, and as shown in the results, was also potentially more powerful in detecting weaker, more latent forms of signals.

To reveal networks associated with deprioritization and prioritization of tasks, firstly, single-trial beta values during ITIs were used to decode active traces of prioritized and unprioritized rules in the previous trials, which yielded seed regions for a subsequent analysis that explored brain regions involved in switching on or off task rules. Specifically, we reasoned that when the priority of a rule was altered, its representational strength between two trials next to each other in areas underlying the shifting process should be correlated. To this end, trial-wise decoding strength of the second trials (Delay 1) for each participant (except for the first trial of each block) were sorted into five trial bins based on the decoding strength within every seed ROI using ITI activity patterns in the first trials. Note that all voxels within the seed regions were used. We tested whether there was a positive linear trend across trial bins by computing Spearman’s correlation between the within-bin averaged strengths and the bin orders (1-5). In essence, this assessed changes in representations in the following trials as a function of the preceding ones^72^. To give an example, to study switching “off” a rule (i.e., from prioritized to unprioritized), trial-wise unprioritized representations were grouped according to the previous trials’ prioritized representations (note that it was the same item, just with different states due to the switch). This was done separately for switch and nonswitch conditions and the resulting coefficients were subtracted from one another (switch minus nonswitch) to identify switch-specific regions with nonswitch trials serving as the natural baseline.

#### Sustained maintenance of unprioritized rules across trials

Following the observation that medial-DMN-centered deprioritization network, which maintained unprioritized rules after a previous switch, was spatially distinct from that of the active unprioritized representations, we reasoned that the separation could be related to the diverging task demands of reducing interference from the uncued task on ongoing trials and sustaining it for later use. To understand whether the latter was supported by the deprioritization network and especially medial DMN, we tested if unprioritized rules were maintained across adjacent trials when they remained uncued by computing between-trial correlations of decoding strengths, focusing on the medial DMN regions as seeds. This followed essentially the same logic as the priority-switching analysis, but instead examining the nonswitch condition, contrasted by the switch condition.

#### Behavioral relevance of trial-wise and block-wise representational strengths

We conducted a series of multiple regression analyses to assess the relationship between the decoding strengths of rules in both priority states and memory performance at different timescales.

For the relation between block-wise decoding strength and behavior, we examined the assumption that the averaged quality of a rule over the course of a block when it was prioritized as well as when it was unprioritized, should similarly facilitate performance on trials when the rule was prioritized, because maintaining both at a longer timescale was essential in a flexible switching paradigm. To test effects of unprioritized rules, we averaged representational strength of the unprioritized rules within a block to predict behavioral measure (sum of zscored absolute color and size errors) of trials in which the respective item became prioritized. That is, for a given trial using rule A, we calculated the mean strength across all trials within the same block when A was unprioritized (Figure 6A) ^41^. The same logic applies when testing effect of the prioritized rule: the averaged prioritized representations across all trials on which A was prioritized, excluding the current trial, was taken as the regressor value. Furthermore, the linear regression model was first fitted at the individual level using block-averaged decoding strength (independently for prioritized and unprioritized rules), with trial numbers within a block and block numbers as additional nuisance predictors to partial out potential influences from order or time. Group-level statistics were calculated by testing individual whole-brain coefficient maps against zero and thresholded using cluster-based permutations. Finally, the unprioritized behavioral results were masked by the deprioritization-specific regions using parcellation-based method (described below) to ensure that the identified regions contained sustained representation of unprioritized rules within deprioritization network. The prioritized behavioral map was similarly intersected by the prioritization network.

For behavioral correlation of trial-wise representations, the models were similar to those of the block-wise variant but instead using the trial-by-trial decoding strengths as predictors. Consistent with previous work ^41^, the decoding strengths were normalized by subtracting the block average to separate variances across trials within a block from overall changes across blocks.

#### Parcellation-based intersection of two statistical maps

As indicated, certain whole-brain results were further masked by another statistical map (Figure 3A, Figure 6BC, and Supplementary Figure 3), using a parcellation-based method. If clusters from the to-be-masked statistical map and the masking map intersected substantially (voxel number >= 30) with the same anatomical parcel defined by the Human Connectome Project brain parcellation (HCP) ^73^, they were characterized as spatially intersecting, and clusters from the to-be-masked map were retained.

## Supplementary figures and tables

**Supplementary Figure 1.**
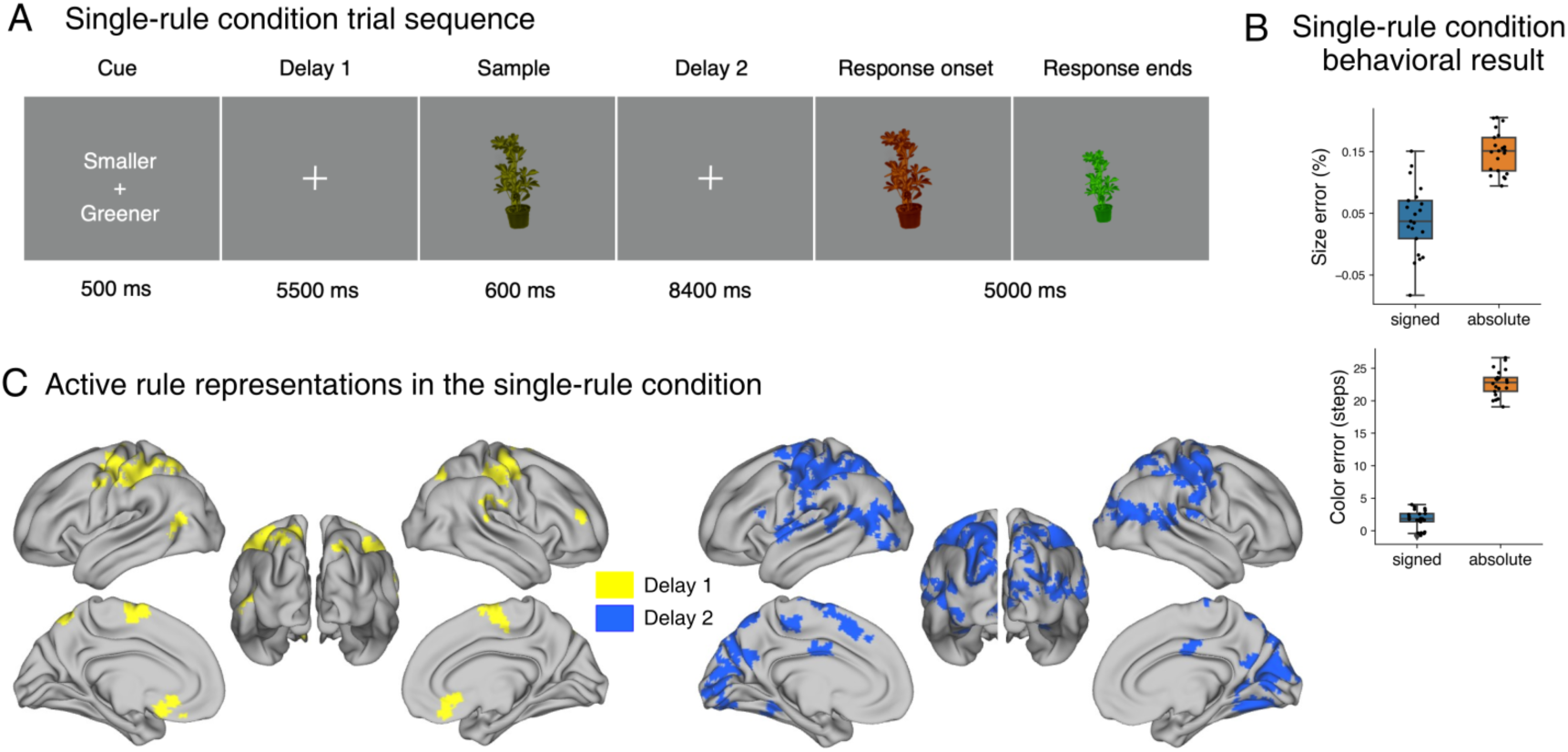
**(A)** Trial sequence for a baseline condition in which task rules were cued on a trial-by-trial basis by texts. Twenty-one of the participants completed this session in addition to the main task on a separate day. **(B)** Size (upper) and color (lower) response errors for the single-rule condition. **(C)** Active decoding of task rules in the single-rule condition. A cluster size of 80 voxels were applied for visualization.

**Supplementary Figure 2.**
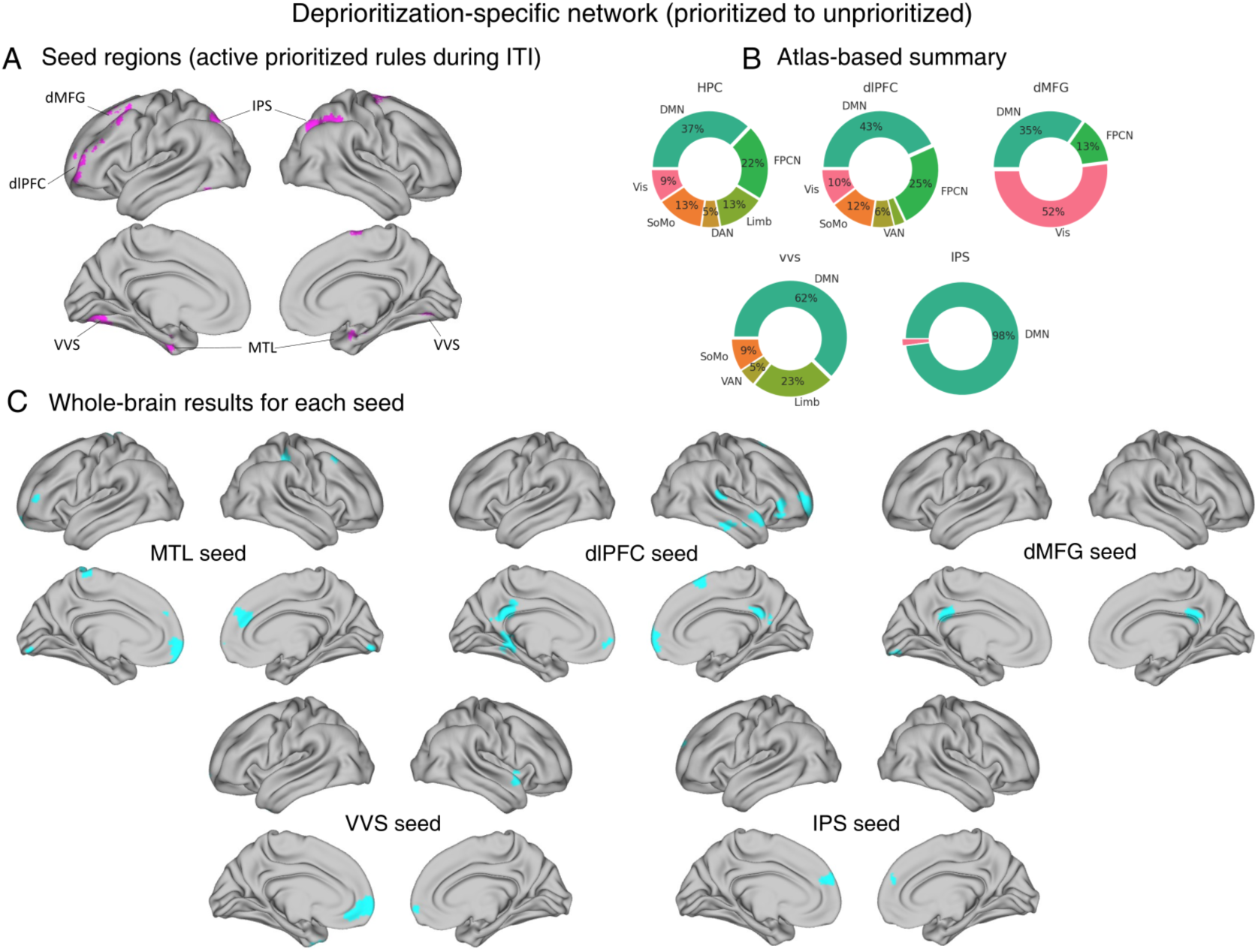
**(A)** Seed regions used in the analysis of deprioritization-specific network. **(B)** Atlas-based summary of the whole-brain maps, separately for each seeded map. **(C)** Significant searchlight results of between-trial correlations (switch – nonswitch) for each seed regions. dMFG = dorsal middle frontal gyrus; dlPFC = dorsolateral prefrontal cortex; IPS = intraparietal sulcus; VVS = ventral visual stream; MTL = medial temporal lobe.

**Supplementary Figure 3.**
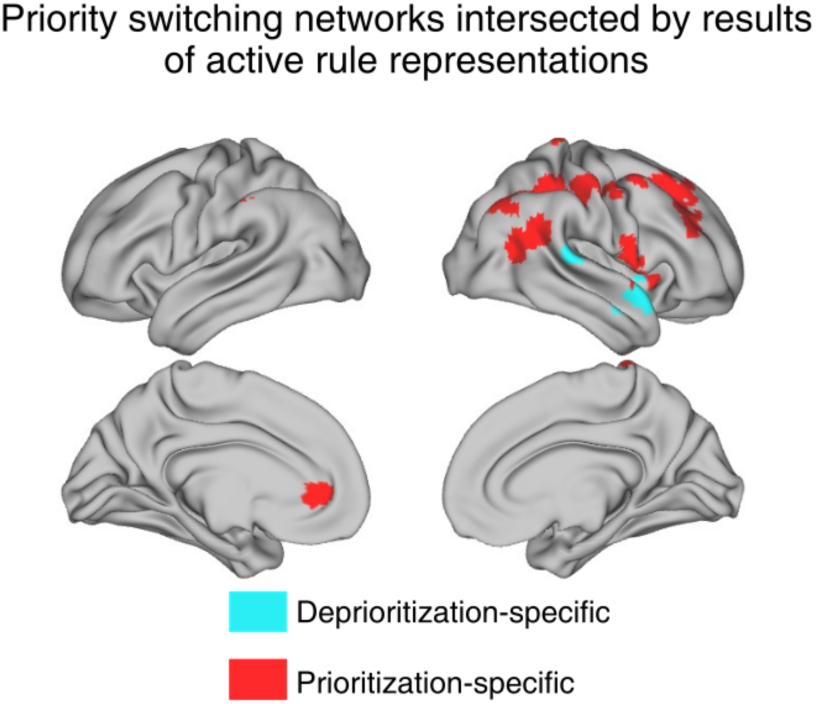
Parcellation-based intersection between the priority switching and active rule representation regions in Figure 2. Cyan: intersection between the deprioritization network and unprioritized representations; Red: intersection between the prioritization network and prioritized representations.

**Supplementary Figure 4.**
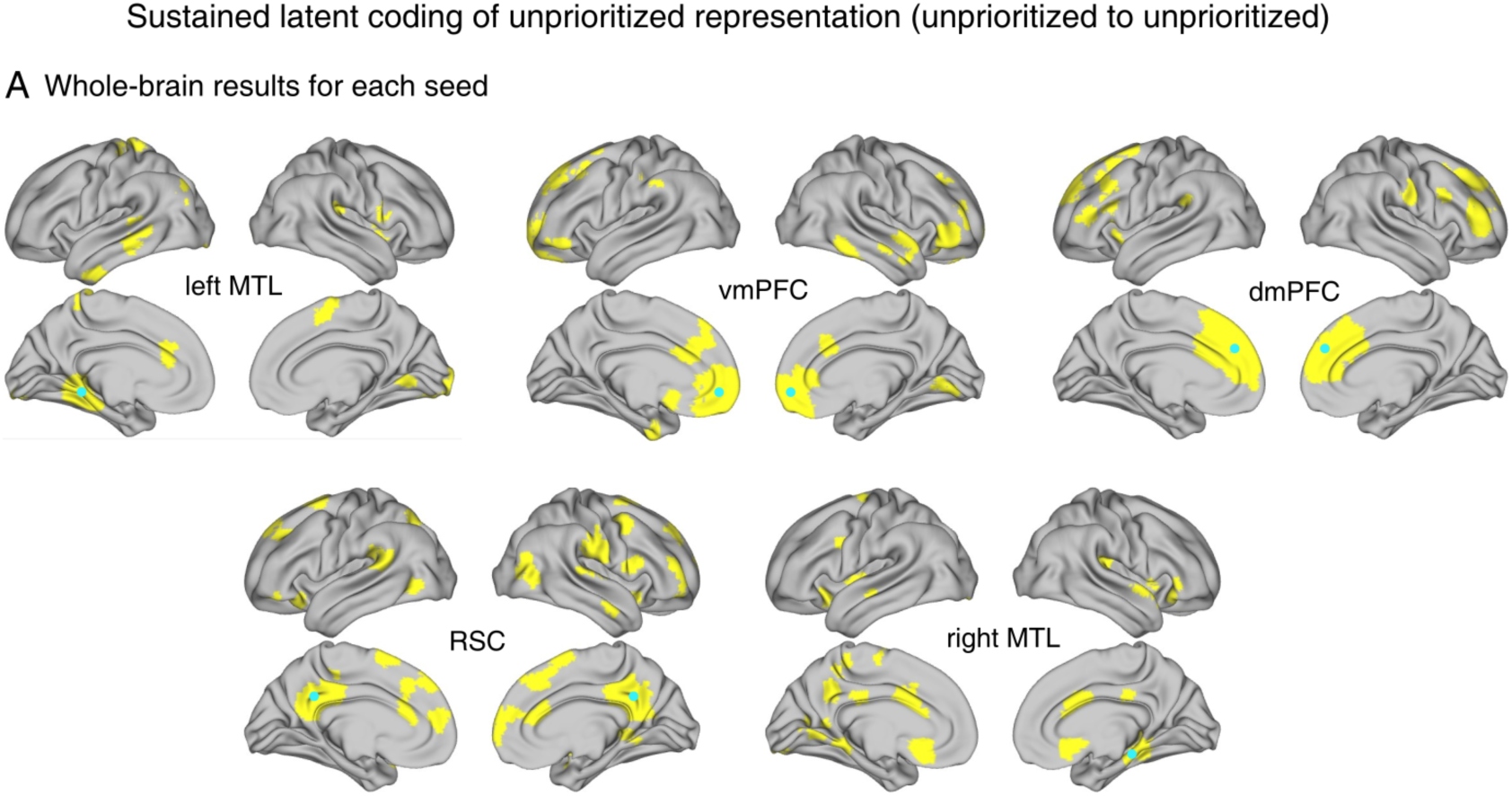
**(A)** Separate maps of sustained unprioritized representations in a latent coding scheme for each seed. Blue dots denote location of the seeds. MTL = medial temporal lobe; vmPFC = ventromedial prefrontal cortex; dmPFC = dorsomedial prefrontal cortex; RSC = retrosplenial cortex.

**Supplementary Figure 5.**
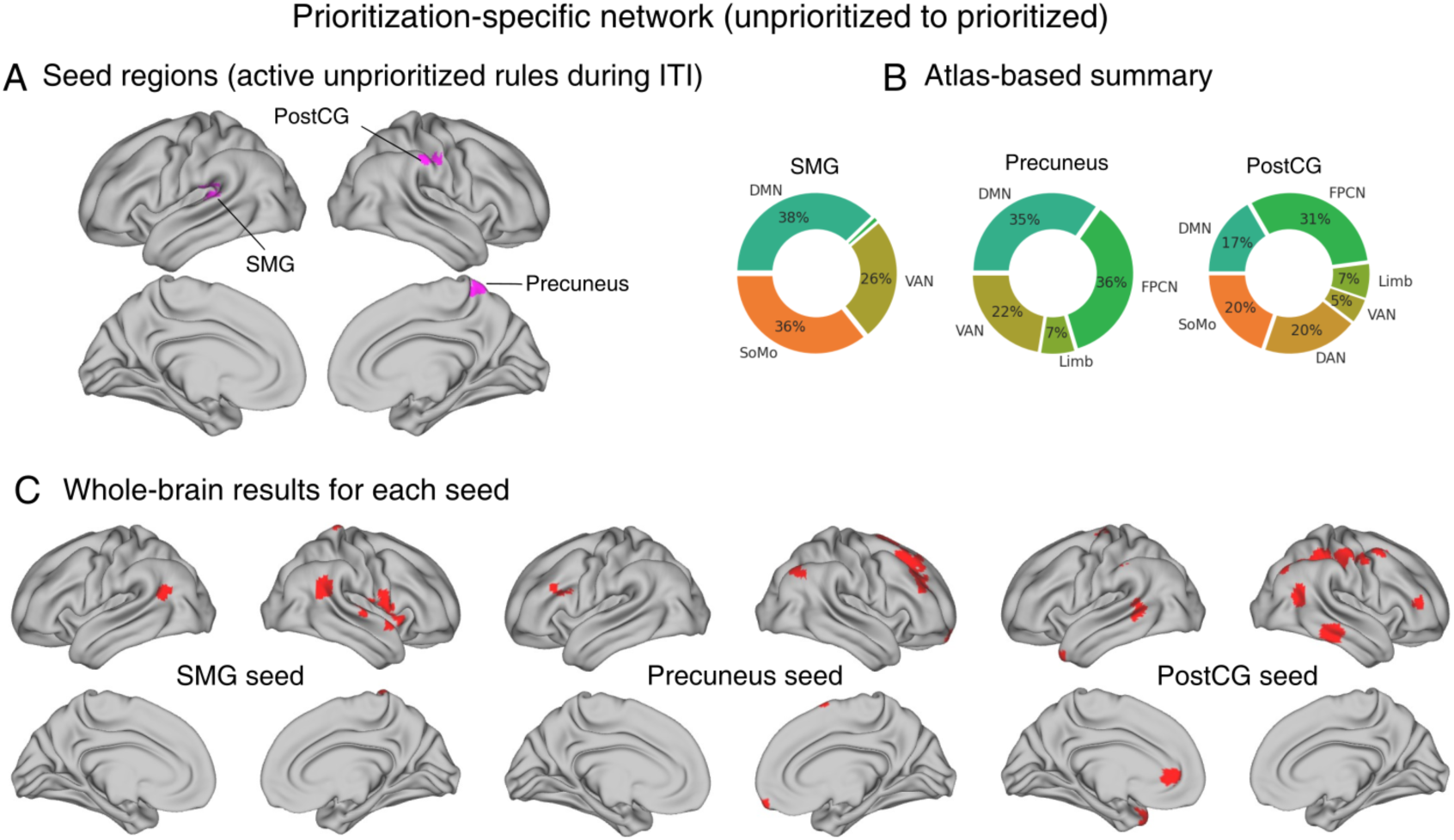
**(A)** Seed regions used in the analysis of prioritization-specific network. **(B)** Atlas-based summary of the whole-brain correlation maps, separately for each seeded map. **(C)** Significant searchlight between-trial correlations (switch – nonswitch) for each seed regions. SMG = supramarginal gyrus; PostCG = postcentral gyrus.

**Supplementary Figure 6.**
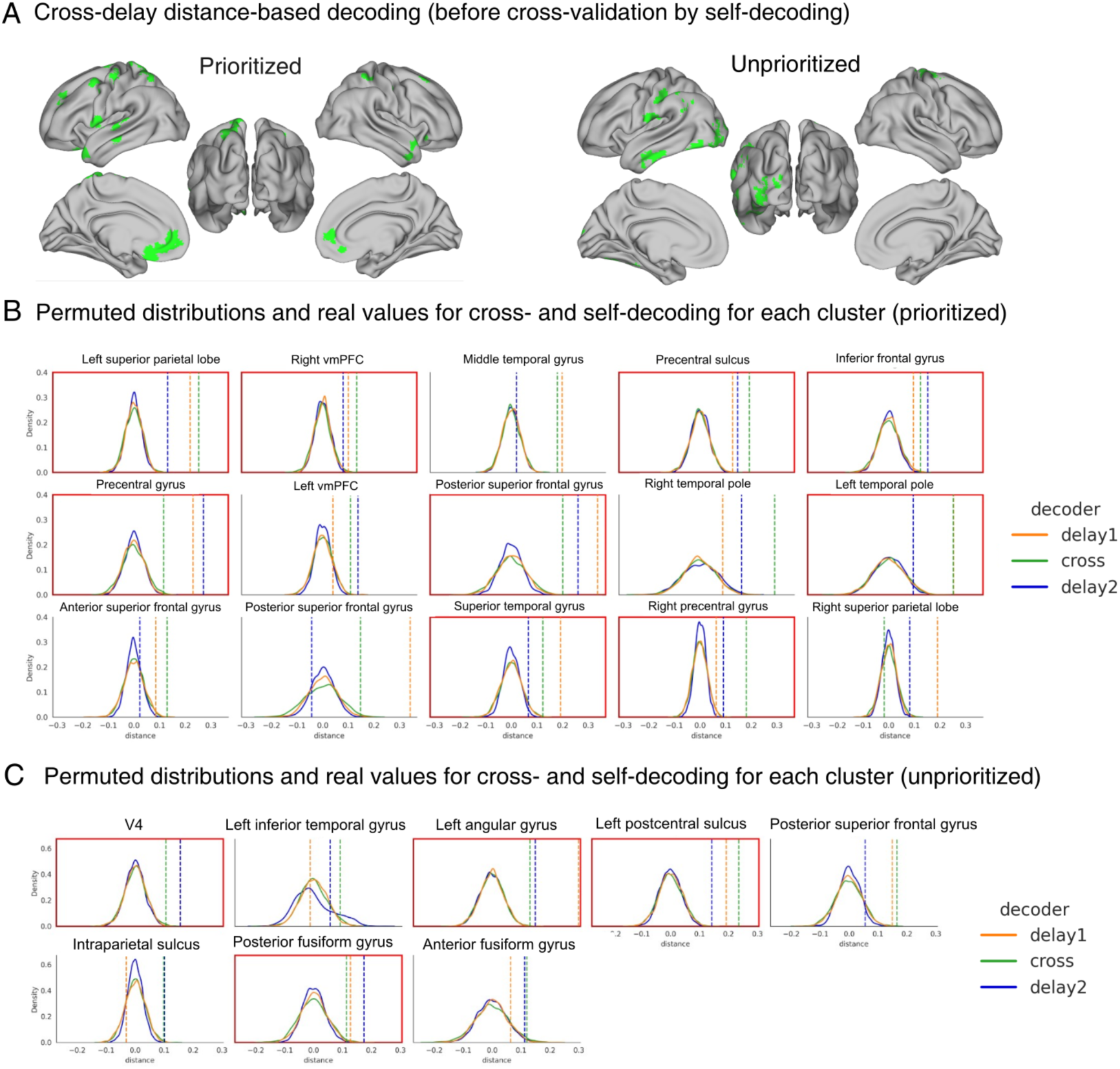
**(A)** Significant clusters showing cross-delay generalization, before cross-validated by distance-based self-decoding (i.e., trained and tested on data from the same delay period). **(B)** Within each cluster from **(A)**, group-level statistics were compared to null distribution created by shuffling trial labels of test sets. Only clusters in which all three decoders (delay1, delay2 and cross-delay) showed significant results (α = 0.05) were included in the final results, as denoted by the red frames surrounding subplots. **(C)** Same as **(B)** but for unprioritized rules.

**Supplementary Table 1.**
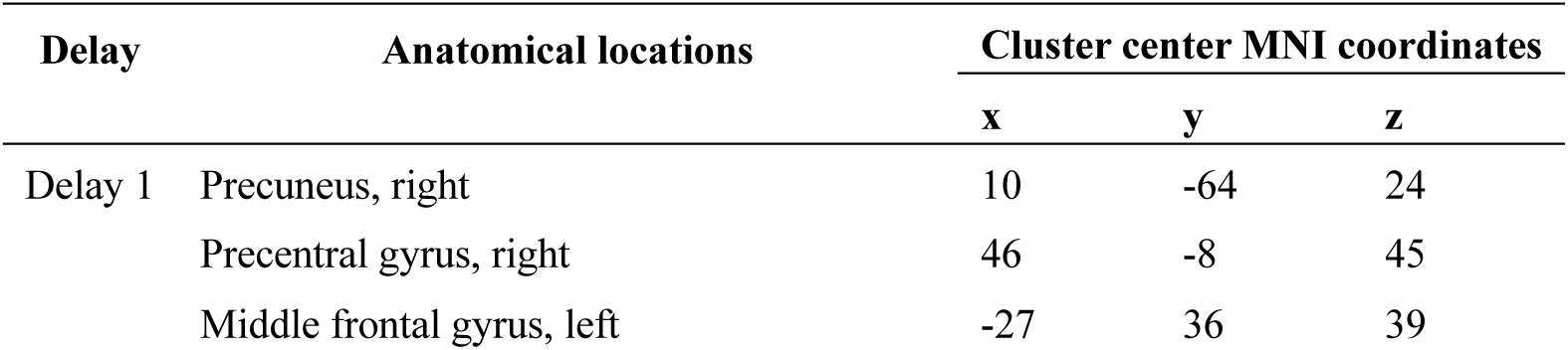
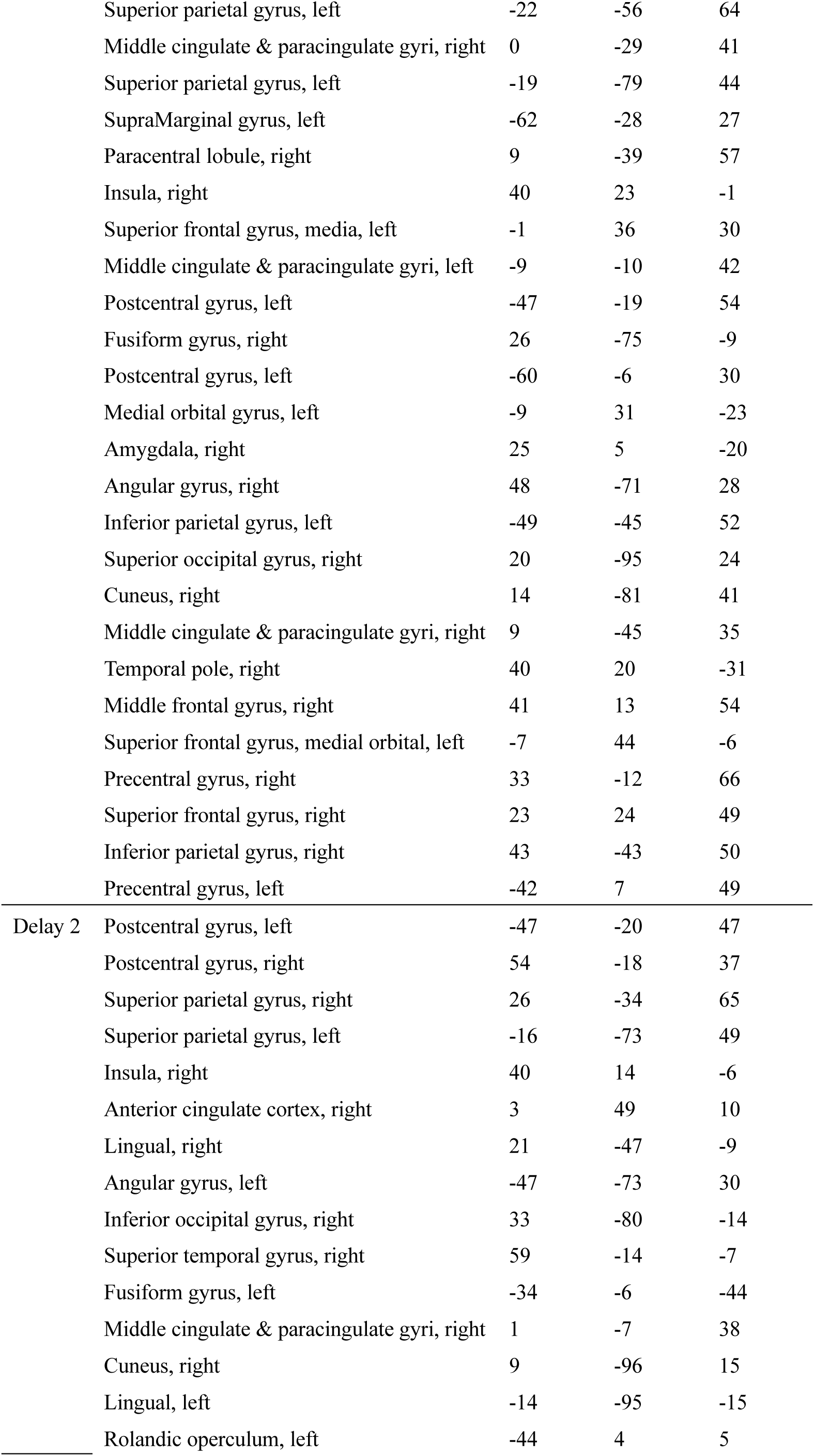
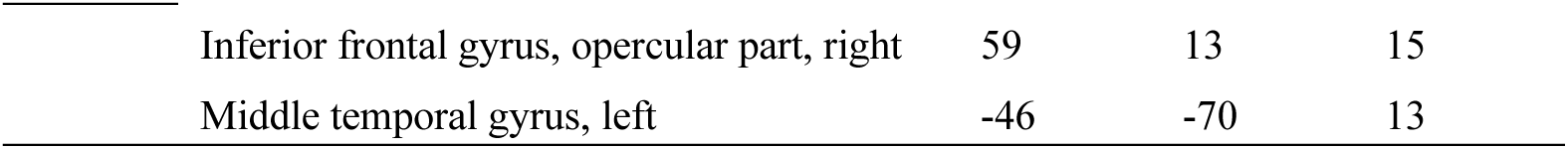
Active representation of prioritized rules.

| Delay | Anatomical locations | Cluster center MNI coordinates |  |  |
| --- | --- | --- | --- | --- |
|  |  | x | y | z |
| Delay 1 | Precuneus, right | 10 | -64 | 24 |
|  | Precentral gyrus, right | 46 | -8 | 45 |
|  | Middle frontal gyrus, left | -27 | 36 | 39 |
|  | Superior parietal gyrus, left | -22 | -56 | 64 |
|  | Middle cingulate & paracingulate gyri, right | 0 | -29 | 41 |
|  | Superior parietal gyrus, left | -19 | -79 | 44 |
|  | SupraMarginal gyrus, left | -62 | -28 | 27 |
|  | Paracentral lobule, right | 9 | -39 | 57 |
|  | Insula, right | 40 | 23 | -1 |
|  | Superior frontal gyrus, media, left | -1 | 36 | 30 |
|  | Middle cingulate & paracingulate gyri, left | -9 | -10 | 42 |
|  | Postcentral gyrus, left | -47 | -19 | 54 |
|  | Fusiform gyrus, right | 26 | -75 | -9 |
|  | Postcentral gyrus, left | -60 | -6 | 30 |
|  | Medial orbital gyrus, left | -9 | 31 | -23 |
|  | Amygdala, right | 25 | 5 | -20 |
|  | Angular gyrus, right | 48 | -71 | 28 |
|  | Inferior parietal gyrus, left | -49 | -45 | 52 |
|  | Superior occipital gyrus, right | 20 | -95 | 24 |
|  | Cuneus, right | 14 | -81 | 41 |
|  | Middle cingulate & paracingulate gyri, right | 9 | -45 | 35 |
|  | Temporal pole, right | 40 | 20 | -31 |
|  | Middle frontal gyrus, right | 41 | 13 | 54 |
|  | Superior frontal gyrus, medial orbital, left | -7 | 44 | -6 |
|  | Precentral gyrus, right | 33 | -12 | 66 |
|  | Superior frontal gyrus, right | 23 | 24 | 49 |
|  | Inferior parietal gyrus, right | 43 | -43 | 50 |
|  | Precentral gyrus, left | -42 | 7 | 49 |
| Delay 2 | Postcentral gyrus, left | -47 | -20 | 47 |
|  | Postcentral gyrus, right | 54 | -18 | 37 |
|  | Superior parietal gyrus, right | 26 | -34 | 65 |
|  | Superior parietal gyrus, left | -16 | -73 | 49 |
|  | Insula, right | 40 | 14 | -6 |
|  | Anterior cingulate cortex, right | 3 | 49 | 10 |
|  | Lingual, right | 21 | -47 | -9 |
|  | Angular gyrus, left | -47 | -73 | 30 |
|  | Inferior occipital gyrus, right | 33 | -80 | -14 |
|  | Superior temporal gyrus, right | 59 | -14 | -7 |
|  | Fusiform gyrus, left | -34 | -6 | -44 |
|  | Middle cingulate & paracingulate gyri, right | 1 | -7 | 38 |
|  | Cuneus, right | 9 | -96 | 15 |
|  | Lingual, left | -14 | -95 | -15 |
|  | Rolandic operculum, left | -44 | 4 | 5 |

|  |  |  |  |
| --- | --- | --- | --- |
| Inferior frontal gyrus, opercular part, right | 59 | 13 | 15 |
| Middle temporal gyrus, left | -46 | -70 | 13 |

**Supplementary Table 2.** Active representation of unprioritized rules.

| Delay | Anatomical locations | Cluster center MNI coordinates |  |  |
| --- | --- | --- | --- | --- |
|  |  | x | y | z |
| Delay 1 | Superior temporal gyrus, right | 65 | -33 | 10 |
|  | Postcentral gyrus, left | -41 | -19 | 49 |
|  | Inferior temporal gyrus, left | -59 | -26 | -24 |
|  | Superior temporal gyrus, left | -64 | -51 | 19 |
|  | Superior occipital gyrus, left | -14 | -75 | 26 |
|  | Fusiform gyrus, left | -37 | -79 | -17 |
|  | Inferior temporal gyrus, right | 50 | -6 | -37 |
|  | Middle occipital gyrus, left | -34 | -88 | 16 |
|  | Fusiform gyrus, left | -27 | -64 | -10 |
|  | Middle temporal gyrus, left | -52 | -72 | 19 |
| Delay 2 | Precentral gyrus, left | -19 | -9 | 69 |
|  | Inferior temporal gyrus, left | -50 | -27 | -28 |
|  | Middle occipital gyrus, left | -21 | -101 | 6 |
|  | Precentral gyrus, right | 37 | -21 | 55 |
|  | Middle occipital gyrus, left | -32 | -90 | 18 |

**Supplementary Table 3.** Deprioritization and prioritization networks.

| Switching<br>direction | Anatomical locations | Cluster center MNI |  |  |
| --- | --- | --- | --- | --- |
|  |  | x | y | z |
| Deprioritization | Superior frontal gyrus, medial orbital, left | -4 | 59 | -6 |
|  | Superior frontal gyrus, right | 26 | 61 | 0 |
|  | Superior frontal gyrus, medial, left | 1 | 48 | 23 |
|  | Temporal pole, right | 54 | 10 | -12 |
|  | Posterior cingulate cortex, left | -0 | -42 | 27 |
|  | IFG pars orbitalis, right | 53 | 33 | -3 |
|  | Parahippocampal gyrus, left | -15 | -38 | -7 |
|  | Lingual, left | -5 | -89 | -13 |
|  | Superior temporal gyrus, right | 68 | -31 | 10 |
|  | Temporal pole, left | -28 | 8 | -43 |
|  | Hippocampus, right | 28 | -34 | -6 |
|  | Supplementary motor area, right | 6 | 16 | 61 |
|  | Paracentral, lobule, left | -6 | -31 | 74 |
|  | Middle frontal gyrus, right | 40 | 17 | 53 |
|  | Inferior temporal gyrus, right | 63 | -21 | -23 |
| Prioritization | Middle frontal gyrus, right | 30 | 29 | 42 |
|  | Angular gyrus, right | 39 | -55 | 44 |
|  | Middle temporal gyrus, right | 50 | -52 | 20 |
|  | Postcentral gyrus, right | 51 | -15 | 47 |
|  | Inferior temporal gyrus, right | 62 | -31 | -17 |
|  | Rolandic operculum, right | 54 | 8 | 2 |
|  | Temporal pole, left | -33 | 17 | -36 |
|  | Middle temporal gyrus, left | -43 | -56 | 19 |
|  | Supplementary motor area, right | 13 | 9 | 67 |
|  | Anterior orbital gyrus, right | 11 | 64 | -17 |
|  | Middle temporal gyrus, left | -56 | -45 | 4 |
|  | SupraMarginal gyrus, left | -64 | -31 | 40 |
|  | Anterior cingulate cortex, left | -4 | 40 | 1 |
|  | Postcentral gyrus, right | 12 | -36 | 79 |
|  | Inferior frontal gyrus, triangular part, left | -49 | 20 | 24 |
|  | Superior temporal gyrus, right | 62 | -13 | 8 |
|  | Insula, right | 39 | -12 | -3 |
|  | Middle frontal gyrus, right | 48 | 41 | 10 |
|  | Precentral gyrus, left | -23 | -16 | 72 |

